# Sculpting the tumour microenvironment by combining radiotherapy and ATR inhibition for curative-intent anti-PD-L1- and anti-NKG2A-based adjuvant immunotherapy

**DOI:** 10.1101/2023.11.08.566202

**Authors:** Emmanuel C Patin, Pablo Nenclares, Charleen Chan Wah Hak, Magnus T Dillon, Anton Patrikeev, Martin McLaughlin, Lorna Grove, Shane Foo, Heba Soliman, Joao P Barata, Joanna Marsden, Holly Baldock, Victoria Roulstone, Joan Kyula, Amy Burley, Lisa C Hubbard, Malin Pedersen, Simon A Smith, Eleanor Clancy-Thompson, Alan A Melcher, Masahiro Ono, Antonio Rullan, Kevin J Harrington

**Affiliations:** Targeted Therapy Team, The Institute of Cancer Research, London, UK; The Royal Marsden Hospital, London, UK; Translational Immunotherapy Team, The Institute of Cancer Research, London, UK; Oncology R&D, Early Oncology Clinical Development, AstraZeneca, Cambridge, UK; Faculty of Natural Sciences, Department of Life Sciences, Imperial College London, UK

## Abstract

Despite some success in other cancer types, the results of combining radiotherapy/chemoradiotherapy and immune checkpoint blockade have been disappointing in patients with locally advanced head and neck squamous cell carcinoma (HNSCC). For such a potentially radiocurable disease, there remains an imperative to explore novel combination approaches. Here, we show that combining ATR inhibition with radiotherapy (ATRi/RT) increases the frequency of highly activated NKG2A/PD-1 double-positive T cells in patients and in animal models of HNSCC. Addition of dual anti-NKG2A and anti-PD-1/-PD-L1 blockade to ATRi/RT in the adjuvant, post-radiotherapy setting induces a robust antitumour immune response in HNSCC preclinical models. Efficacy of the combination regimen relies on CD40/CD40L costimulatory-mediated infiltration of activated/proliferative/memory CD8 and CD4 conventional T cells with persistent or new T cell receptor (TCR) signalling, respectively, as defined by tracking of T cell dynamics. In this favourable therapeutic context, TCR sequencing shows increased richness of the TCR repertoire and the emergence of numerous and large TCR clusters that share antigen specificity in response to full combination therapy. Collectively, our data point towards promising combination approaches for future clinical testing in HNSCC.

## Introduction

New treatment modalities are urgently needed to reduce the number of patients that relapse after multimodality treatment for locally advanced head and neck squamous cell carcinoma (HNSCC). To achieve this goal, we must improve our understanding of strategies that seek to combine radiotherapy, chemotherapy, targeted therapy and/or immunotherapy (ImmuT). Immune checkpoint blockade (ICB) has significant activity as a single-agent and in combination with chemotherapy in recurrent/metastatic HNSCC^1, 2, 3^. However, concomitant addition of ICB has so far shown no evidence of improving standard-of-care radiotherapy/chemoradiotherapy for locally advanced HNSCC^4, 5^ (NCT03040999). Since simple addition of concomitant anti-PD1/-PD-L1 blocking antibodies to radiotherapy/chemoradiotherapy fails to improve outcomes, alternative approaches involving combinations with antibodies targeting other immune checkpoints, including anti-NKG2A, are under preclinical and clinical development.

The inhibitory receptor NKG2A is a member of the C-type lectin superfamily expressed on both T and NK cells^6^. Downstream inhibitory signalling via NKG2A relies on the recruitment of SHP-1/2 phosphatases which subsequently suppress the activity of activating receptors such as NKG2D. Preclinical data indicated that anti-NKG2A synergised with anti-PD-1/PD-L1 by boosting the activity of NK and CD8 T cells^7^. NKG2A inhibition also delayed tumour growth after peptide vaccination in various mouse tumour models^8^. In clinical studies, the humanised anti-NKG2A antibody monalizumab, in combination with durvalumab, showed manageable toxicities and promising activity in patients with microsatellite-stable colorectal cancer^9^. The same combination, tested in the COAST and NeoCOAST studies, increased therapeutic benefits when compared to durvalumab alone in non-small-cell lung cancer^10, 11^. In the context of HNSCC, monalizumab alone did not significantly improve therapeutic outcome in the UPSTREAM clinical trial^12^. Nevertheless, evaluation of monalizumab, in combination with durvalumab, is under investigation in the same trial. The recent failure of the INTERLINK-1 clinical trial (NCT04590963) of monalizumab and cetuximab in recurrent and/or metastatic HNSCC has refocused efforts on combinations of monalizumab with anti-PD1/-PD-L1 agents.

Another strategy to augment the efficacy of ICB is through combination with RT and radiosensitizers, such as DNA damage response inhibitors (DDRi). One such DDRi, the ATR inhibitor (ATRi), ceralasertib, has been under clinical investigations for head and neck cancer^13^. We and others previously reported that ATRi could potentiate RT-mediated antitumour immunity, creating a favourable tumour microenvironment (TME) for ICB interventions^14, 15, 16^. Indeed, combining ATRi/RT with anti-PD-L1 blockade significantly improved tumour growth control and survival in mice^17, 18^. Additionally, our group also recently described how dual anti-PD-1/-TIGIT blockade could enhance NK-cell-mediated tumour control after ATRi/RT^15^. Following those preclinical studies, combination of ATRi, RT and ICB is now entering clinical trials for the treatment of HNSCC (NCT04576091 and EudraCT 2020-001034-35).

Here, we hypothesised that targeting both NKG2A and PD-1/PD-L1 axes with blocking antibodies would boost the antitumour response following ATRi/RT. By interrogating published single-cell RNAseq datasets, we demonstrate the presence of highly activated NKG2A/PD-1 double-positive T cells in human head and neck tumours. Analyses from our own clinical samples reveal that the frequencies of those T cells can be increased following ATRi/RT, making them an interesting target for immunotherapeutic modulation following combined RT and ATRi therapy. Using HPV^-^ and HPV^+^ preclinical models of head and neck cancer, we describe for the first time how the combination of ATRi, radiation and immunotherapy, based on dual anti-NKG2A and anti-PD-L1 immune checkpoint blockade (ATRi/RT/ImmuT), induces a robust antitumour response. Treatment efficiency is dependent on the recruitment of both CD8 and CD4 conventional T cells which display highly activated, proliferative and cytotoxic phenotypes in the tumour microenvironment (TME). Blocking antibody studies reveal that activation and proliferation of both CD8 and CD4_conv_ T cells is dependent on the CD40/CD40L signalling pathway. To understand further the therapeutic effect of the combination, we also report that addition of immunotherapy to ATRi/RT significantly modulates the temporal dynamics of T cell activation and influences TCR repertoire richness, clonality, and antigen specificity in the TME. Together, these data present a strong case for rational clinical trial designs incorporating ATRi/RT followed by adjuvant doublet immunotherapy targeting NKG2A and PD-1/PD-L1.

## Results

### Highly activated NKG2A/PD-1 double positive T cells are present in the tumour microenvironment of HNSCC patients

The combination of anti-NKG2A and anti-PD-1/PD-L1 axis ICB has recently gained momentum in clinical settings, particularly in lung cancer. Reinvigoration of NKG2A/PD-1 double-positive cytotoxic lymphocyte cells in the tumour microenvironment (TME) could provide a mechanistic basis for the combination, including in HNSCC. Thus, we interrogated published databases in patients with HNSCC. First, analysis of the Cancer Genome Atlas (TCGA) revealed a significant positive correlation between expression of *KLRC1* (NKG2A) and both *PDCD1* (PD-1) and *CD274* (PD-L1) in HPV^-^ and HPV^+^ HNSCC tumours (figure 1A). To identify the type of immune cells expressing both *KLRC1* and *PDCD1*, we analysed single-cell RNA sequencing (scRNAseq) databases available in two previously published articles^19, 20^. Interestingly, *KLRC1/PDCD1* double-positive immune cells were detected in both studies and those cells were predominantly present in tumours rather than blood (figure 1B and S1A). Further detailed analyses also showed that the majority of *KLRC1*+/*PDCD1*+ cells were in the CD8^+^ T cell compartment (figure 1C-D and figure S1B-C). Additionally, we observed that *KLRC1/PDCD1* double-positive CD8 T cells were highly activated, as shown by high expression of cytotoxic/effector/memory/proliferating lymphocyte gene signatures including *GZMA*, *GZMB*, *PRF-1*, *IFNG*, *CCL5*, *TIGIT*, *LAG3*, *ITGAE*, *CD69* and *PCNA* (figure 1E-F and figure S1D-E).

**Figure 1.**
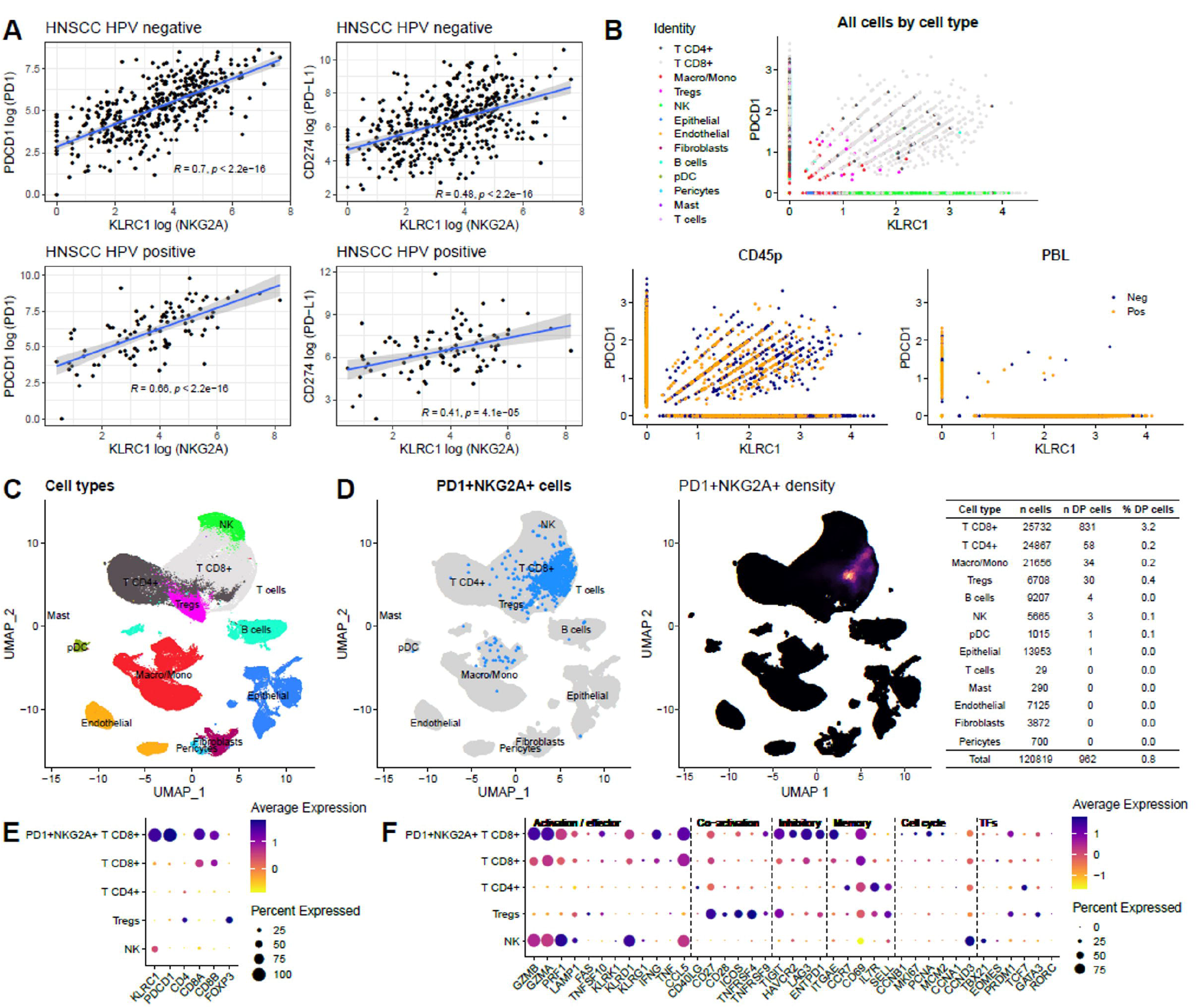
Highly activated NKG2A and PD-1 double-positive T cells are detected in patients with head and neck cancer patients. (A) TCGA database analyses of the correlation between expression of *KLRC1* and both *PDCD1* and *CD274* in both HPV^-^ and HPV^+^ HNSCC patients. (B) Scatter plots showing expression of *KLRC1* and *PDCD1* in various cell populations (higher panel) and tumours versus blood (lower panels) (C) UMAP plot showing distribution of identified cell clusters distribution. (D) Left panel; UMAP plot depicting NKG2A/PD-1 double positive (DP) cells in the different identified cell clusters, right panel; UMAP plot depicting intensity expression of NKG2A/PD-1 double positive cells with absolute numbers for each identified cell populations (right table). Dot plots showing average and percentage expression of the *KLRC1*, *PDCD1* and pan-cell markers (E) and activation/effector, co-activation, inhibitory, memory, cell cycle and transcription factors (TFs) markers (F) in NKG2A^+^/PD-1^+^ CD8^+^ T cells versus CD8^+^T, CD4^+^T, Tregs and NK cells. Datasets from Kurten CHL et al., 2021.

Taken together, these analyses of previously published scRNAseq datasets show the presence of highly activated NKG2A/PD-1 double-positive CD8 T cells in the TME of patients with HNSCC and point towards opportunities for their clinical exploitation.

### Combining ATR inhibition and RT increases the frequencies of NKG2A/PD-1 double-positive CD8 and CD4 T cells in patients with HNSCC

Because of the high activation status of NKG2A/PD-1 double-positive T cells and their potential as therapeutic targets, we tested whether combination of ATRi and RT increased the number of these cells in patients. Thus, we analysed surface expression of both NKG2A and PD-1 in PBMCs of six head and neck cancer patients enrolled in the phase I PATRIOT clinical trial (NCT02223923)^21^. In this trial, patients were treated with an ATRi (ceralasertib, AZD6738, 80 mg twice daily) in combination with 5 daily fractions of 2 Gy radiotherapy per week for 3 weeks, starting 3-7 days after commencing ceralasertib. Blood samples were collected at 4 time-points: baseline, pre-RT fraction 1 (pre-F1), pre-RT fraction 6 (pre-F6) and pre-RT fraction 11 (pre-F11). Over time, we observed that the ATRi and RT combination treatment increased the subpopulation of NKG2A/PD-1 double-positive CD8 but also, interestingly, CD4 T cells (figure 2A-B). Notably, increased proportions of both CD8 and CD4 T cells were detected in both partial (PR) and complete (CR) responders, but not in patients achieving a best response of stable disease (SD) (figure 2A-B).

**Figure 2.**
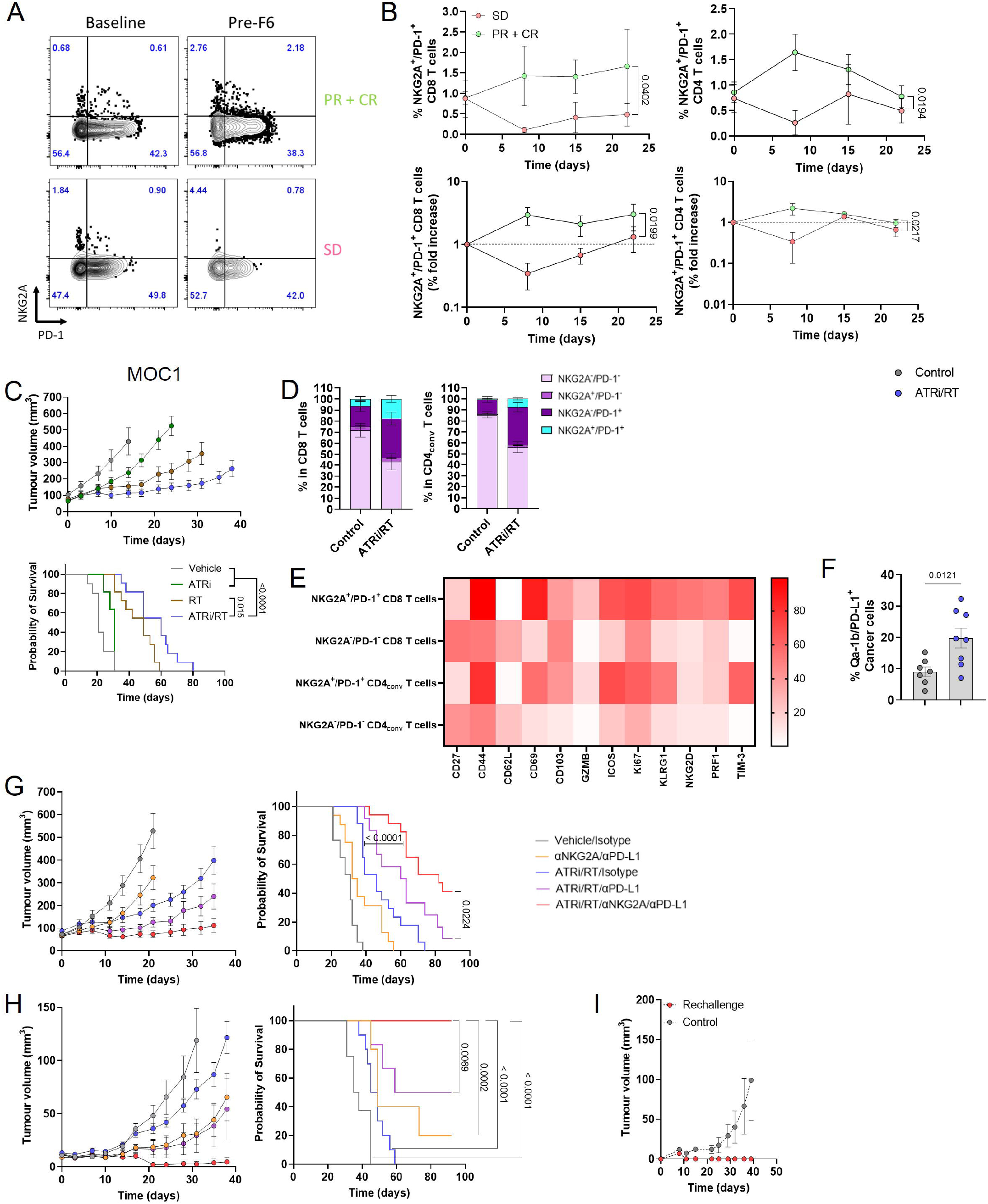
NKG2A and PD-1/PD-L1 axis immune checkpoint blockade improves the therapeutic outcome of ATRi/RT. (A) Dot plots showing NKG2A and PD-1 surface expression on CD8 T cells in one patient from the partial/complete response (PR + CR) group versus one patient from the stable disease response (SD) group. (B) % and fold increase of NKG2A/PD-1 double positive CD8 and CD4 T cells in PR + CR versus SD group (3 patients/group). Experiments in this figure were performed in the MOC1 model. (C) Tumour growth and survival curves across the different conditions (10-11 mice/group). (D) Bar chart; % of NKG2A and/or PD-1 positive populations in CD8 and CD4_conv_ T cells, scatter plot with bar; NKG2A/PD-1 positive populations in CD8 and CD4_conv_ T cells (8-12 mice/group). (E) Heatmap showing marker intensity of expression in NKG2A^+^/PD-1^+^ versus NKG2A^-^/PD-1^-^ CD8 and CD4_conv_ T cells. (F) % surface expression of Qa-1b/PD-L1 double positive cancer cells in the different conditions. Tumour growth and survival curves across the different conditions in ectopic (G) and orthotopic (H) (5-17 mice/group). (I) Tumour growth in control versus rechallenged mice (5-6 mice/group).

Therefore, the detection of both NKG2A/PD-1 double-positive CD8 and CD4 T cells in patients enrolled in the PATRIOT clinical trial, particularly in good responders, provides a rationale for the combination of both anti-NKG2A and anti-PD-1/-PD-L1 axis immune checkpoint blockades (ImmuT) following chemoradiotherapy.

### Dual anti-NKG2A and anti-PD-L1 immune checkpoint blockade enhances ATRi/RT-mediated antitumour response in head and neck cancer preclinical models

Before evaluating the hypothesis that a combination of anti-NKG2A and anti-PD-1/-PD-L1 targeted antibodies could act in concert with ATRi/RT to improve tumour control, we quantified NKG2A/PD-1 double-positive T cells in the TME following ATRi/RT treatment in both HPV^-^ (MOC1) and HPV^+^ (mEER) preclinical models^22, 23, 24^.

We confirmed that combined ATRi/RT delayed tumour growth and increased survival in both models when compared to single-agent treatment (figure 2C and figure S2A). Critically, ATRi/RT led to an increase of the proportions of both NKG2A/PD-1 double-positive CD8 and CD4 conventional (CD4_conv_) T cells in both MOC1 and mEER ectopic (subcutaneous) tumours (figure 2D and figure S2B-C). Those cells were more activated than their NKG2A/PD-1 double-negative counterparts, as shown by higher expression levels of activation/memory/proliferation markers, such as CD44, CD69, GZMB, PRF-1, Ki67, NKG2D and TIM-3 (figure 2E). Of note, increased proportions of NKG2A/PD-1 double-positive CD8 and CD4_conv_ T cells were accompanied by a greater number of cancer cells expressing both Qa-1b and PD-L1, the ligands for NKG2A and PD-1, respectively (figure 2F and figure S2D). These results strengthen the hypothesis that ATRi/RT can create a TME in which the NKG2A/Qa-1b and PD-1/PDL-1 axes might favourably be targeted concomitantly.

Next, mice bearing ectopic MOC1 or mEER tumours received systemic anti-NKG2A and/or anti-PD-L1 blocking antibodies after ATRi/RT. Interestingly, the combination of both anti-NKG2A and PD-L1 antibodies significantly improved ATRi/RT treatment in terms of tumour growth and survival in both models (figure 2G and figure S2E). The addition of anti-NKG2A was necessary, as anti-PD-L1 antibody treatment alone exerted more modest benefits. Importantly, the actual therapeutic benefit of having the full combination of ATRi/RT/anti-NKG2A/anti-PD-L1 was further confirmed orthotopically using the MOC1 model. Strikingly, when compared with the other treatment groups, all mice treated with the full combination had complete tumour eradication (figure 2H). Finally, previously cured animals did not develop tumours on rechallenge in both models (figure 2I and figure S2F).

Taken together, these data show that ATRi/RT increased the frequencies of both NKG2A^+^/PD-1^+^ CD8 and CD4conv T cells, alongside Qa-1b^+^/PD-L1^+^ cancer cells in the TME. Subsequent adjuvant therapy with NKG2A and PD-L1 blocking antibodies after ATRi/RT stimulated favourable effector and memory antitumour immunity in both HPV- and HPV+ head and neck cancer preclinical models.

### Combination of ATRi/RT and anti-NKG2A/PD-L1 dual immune checkpoint blockade induces gene signatures characteristic of T cell activity

To study the nature of the antitumour immune response following ATRi/RT, we performed RNA sequencing (RNAseq) in MOC1 and mEER tumours on day 11 post-radiation (8 days after the beginning of immunotherapy). Differentially gene expression (DEG) analysis showed significant changes induced by ATRi/RT, which were further increased by adding immunotherapy (figure 3A and figure S3A). Volcano plots also revealed higher expression of genes typical of an immune response, such as *PRF-1*, *CD274* and *CD8a*, when mice were treated with the full combination therapy (figure 3B and figure S3B).

**Figure 3.**
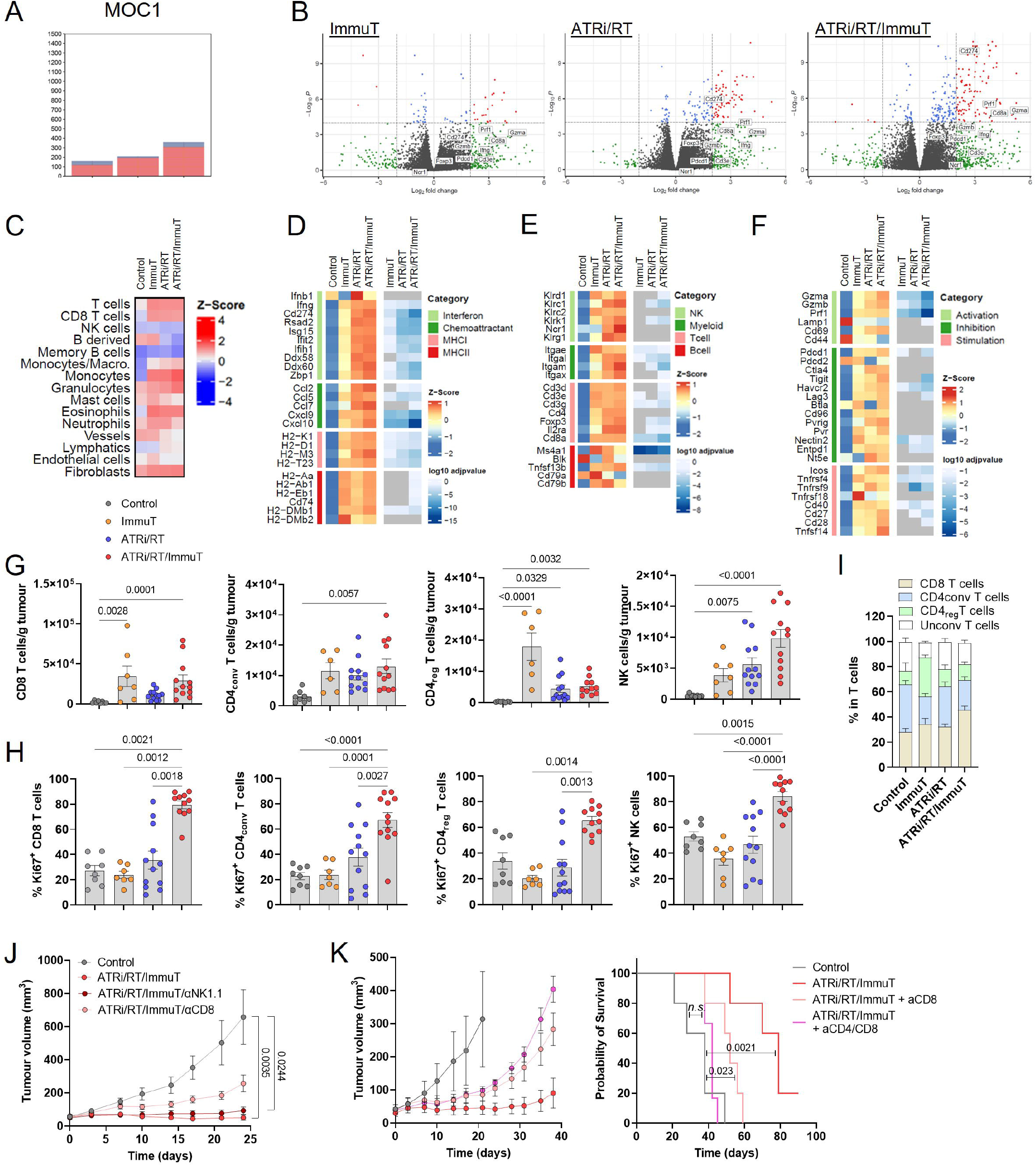
ATRi/RT and anti-NKG2A/PD-L1 immunotherapy trigger a potent T-cell antitumour immune response. Experiments presented in this figure were performed in the MOC1 model. (A) Number of differentially expressed genes (DEGs) across the different conditions calculated by DESeq2. (B) Volcano plots showing expression of the different genes in the various conditions (genes characteristic of an immune response are named). (C) Immune cell population estimates. Immune cell scoring was performed on normalised RNAseq counts using the mMCP-counter package. Heatmaps corresponding to interferon and cytokine signalling, chemoattractant, MHCI and MHCII (D); immune cell populations (E); and immune cell activation status (F). Data shown are z-scores of log2 transformed normalised counts for the treatment conditions shown. This plotted alongside log10 adjusted *p*-value for each gene calculated from DEG analysis using DESeq2. Non-significant adjusted *p*-values > 0.05 are indicated as grey. (G) Absolute number/gram of tumour of the indicated lymphocytes in the various conditions (8-12 mice/group). (H) % Ki67-positive cells in the indicated lymphocytes in the different conditions (8-12 mice group). (I) % indicated lymphocytes in the total T cell population across all treatment conditions. (J) Tumour growth curves across all conditions (5-6 mice/group). (K) Tumour growth and survival curves across the different conditions (5-6 mice/group).

Immune cell population estimates from transcriptomic data, using mMCP-counter, highlighted the prominence of T cells in both models in response to treatment (figure 3C and figure S3C). However, there were some differences between the two models in the case of monocytes, granulocytes and fibroblasts. Analysis of single gene expression showed that, in the MOC1 model, ATRi/RT/anti-NKG2A/anti-PD-L1 (ATRi/RT/ImmuT) combination, when compared with the other groups, regularly induced more interferon signalling (*CD274, RSAD2, ISG15, IFIH1, DDX60* and *ZPD1*), chemoattraction (*CCL2, CCL5, CCL, CCL9* and *CXCL10*), MHCI-(*H2-Aa, H2-D1, H2-M3* and *H2-T23*) and MHCII-related (*H2-AA, H2-AB1, H2-EB1, CD74* and *H2-DMB1*) gene expression (figure 3D). Generally, differences were less pronounced in the mEER model, with the exception of MHCII-related genes (figure S3D). For immune cell population gene signatures, in response to ATRi/RT/ImmuT, the lowest numerical *p*-values were observed for different NK cell receptors, such as *KLRD1, KLRC1* and *KLRK1* in MOC1 tumours but only *KLRD1* and *KLRK1* in mEER tumours (figure 3E and figure S3E). It is remarkable that, in both models, we observed higher increases of genes characteristic of T cell activity, including *CD3D*, *CD3E*, *FOXP3* and *CD8A*, in response to ATRi/RT/ImmuT (figure 3E and figure S3E). Finally, we also detected in the same treatment group high expression of genes for immune cell activation markers, including *GZMA, GZMB, PRF1, PDCD1, HAVCR2, ENTPD1, ICOS, TNFRSF, CD40, CD27* and *CD28*, in both MOC1 and mEER tumours (figure 3F and figure S3F).

Together, RNAseq analysis suggests that ATRi/RT and anti-NKG2A/PD-L1 dual immune checkpoint blockade triggers a robust immune response in the TME, which is likely to be mediated by the activity of T cells in both HPV^-^ and HPV^+^ head and neck cancer models.

### The efficacy of the combination of ATRi/RT and anti-NKG2A/PD-L1 immunotherapy is dependent on the activity of both CD8 and CD4 T cells

To confirm the RNAseq results, we performed flow cytometry analyses in both MOC1 and mEER tumours. The lymphocyte data revealed that full combination therapy with ATRi/RT/ImmuT significantly induced greater infiltration of CD8, CD4_conv_ and CD4 regulatory (reg) T cells, as well as NK cells, into the TME in both models in comparison with the other treatment groups (figure 3G and figure S3G). Of note, ATRi/RT or ImmuT on their own did significantly boost the infiltration of some lymphocyte subtypes, such as CD8 and CD4_conv_ T cells, in some instances. Importantly, however, only the full ATRi/RT/ImmuT combination led to a significant improvement of lymphocyte activation as demonstrated by increased intracellular expression of the pan-activation/proliferation marker Ki67 (figure 3H and figure S3H). In addition, in respect of T cell compartment, ATRi/RT/ImmuT caused increased frequencies of CD8 T cells (figure 3I and figure S3I). Finally, to prove specific reactivity of T cells towards tumour antigens, we restimulated splenocytes from mEER tumour-bearing mice treated with the different therapies with a long overlapping HPV16 E7 peptide pool. Only full combination therapy significantly increased the proportions of both CD8 and CD4_conv_ T cells with proliferative and cytotoxic capacities (figure 3SJ).

In light of those data, we defined the subtypes of lymphocytes mediating treatment efficacy using specific depleting antibodies. In mice treated with ATRi/RT/ImmuT, the acceleration of tumour growth and shortened lifespan in the absence of CD8 T cells demonstrated the crucial role of those cells in mediating therapy in the MOC1 model (figure 3J). The effect of CD8 T cells was even more marked in the mEER model (figure S3K). In contrast to CD8 T cells, however, depletion of NK cells did not have a significant effect on the efficacy of the full combination treatment (figure 3J and figure S3K). Strikingly, the depletion of both CD8 and CD4 T cells in MOC1-bearing mice completely abrogated the effect of the full combination therapy in terms of mouse survival (figure 3K). Delayed separation between the tumour growth curves of CD8 and CD8/CD4 T cell-depleted mice suggests a delayed role for CD4 T cells.

Taken together, these data suggest that the efficacy of ATRi/RT/ImmuT combination therapy is dependent on the infiltration of both activated CD8 and CD4 T cells in the TME.

### Combination of ATRi/RT and anti-NKG2A/-PD-L1 immunotherapy promotes the infiltration of CD8 and CD4_conv_ T cell subpopulations with a PD-1^high^ cytotoxic/proliferative/effector memory phenotype

Next, we sought insight into the specific phenotypes of therapy-related tumour-infiltrating CD8 and CD4 T cells. Thus, we performed extensive immune profiling in the MOC1 model, using a combination of TriMap and FlowSOM algorithms to identify the various clusters of CD8 and CD4 T cell populations in control and treatment groups.

We observed a disappearance of naïve/inactivated CD8 and CD4 T cell populations (pop 4 and 0, respectively) from MOC1 tumours. These populations were diminished in the ATRi/RT cohort and completely absent in response to the full combination therapy (figure 4A-B). In parallel, a shift towards populations with an activated/proliferative phenotype for CD8 T cells was identified in response to ATRi/RT (pop 6 and 7), but this effect was accentuated after the addition of immunotherapy (pop 2 and 3) (figure 4A). The same pattern was detected for CD4_conv_ T cells, with populations displaying more pronounced activated/proliferative phenotypes (pop 3, 4, 6 and 9) in the full combination therapy when compared with ATRi/RT alone. By contrast, adding immunotherapy to ATRi/RT did not significantly modulate the various populations of CD4_reg_ T cells when compared with ATRi/RT alone (figure 4B).

**Figure 4.**
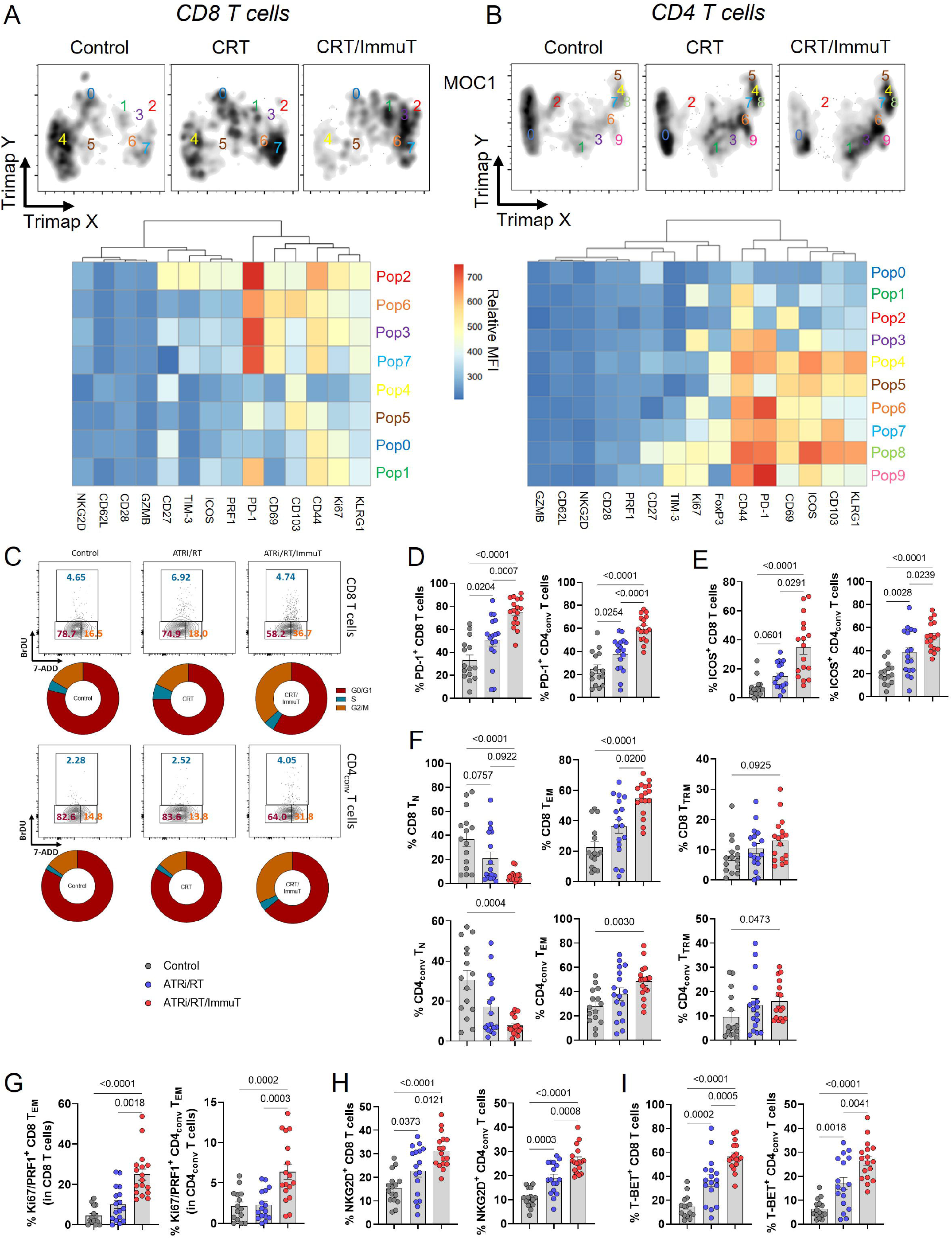
ATRi/RT and anti-NKG2A/PD-L1 immunotherapy induce the proliferation of PD-1^+^ effector memory cytotoxic CD8 and CD4 T-cells in the tumour microenvironment. Experiments presented in this figure were performed in the MOC1 model. (A and B) Higher panels; intensity of the different subpopulation clusters of CD8 (A) and CD4 (B) T cells identified using TriMap and FlowSOM algorithms, lower panels; heatmap showing the expression of the indicated markers in the different populations identified by FlowSOM algorithm across all treatment conditions (concatenated from 7-8 mice/group). (C) Dot plot and donut chart representing % of CD8 and CD4_conv_ T cells in the different cell cycle phases across all conditions (concatenated from 5-6 mice/group). Across all conditions the following graphs show % (D) PD-1^+^, (E) ICOS^+^, (F) naïve (N; CD62L^+^ CD44^-^), effector (EM; CD62L^-^ CD44^+^ CD103^-^) and tissue-resident (TRM; CD62L^-^ CD44^+^ CD103^+^) memory, (G) Ki67/PRF^+^ EM, (H) NKG2D^+^ and (I) T-BET^+^ CD8 and CD4_conv_ T cells (15-17 mice group).

The key findings from TriMap/FlowSOM analyses were further investigated to evaluate their significance. Since increased proliferation was an important feature of T cell subpopulations infiltrating MOC1 tumours after full combination therapy, we tracked cell cycle distribution in CD8 and CD4_conv_ T cells. ImmuT added to ATRi/RT caused a noticeable shift of both CD8 and CD4_conv_ T cells towards G2/M phase, which suggests the presence of actively dividing cells (figure 4C). At the same time, we observed that ATRi/RT/ImmuT was instrumental in increasing the frequencies of PD-1^+^ (figure 4D), ICOS^+^ (figure 4E) and effector memory (EM) (figure 4F) CD8 and CD4_conv_ T cells. In line with the TriMap/FlowSOM analysis, MOC1 tumours were more infiltrated with Ki67/PRF1^+^ CD8 and CD4_conv_ T_EM_ cells in response to the full combination (figure 4G). We also detected elevated frequencies of NKG2D^+^ and T-BET^+^ CD8 and CD4_conv_ T cells in the same treatment group when compared with ATRi/RT alone (figure 4H-I).

To evaluate the specific effect of including anti-NKG2A antibody in full combination therapy at the cellular level, we performed the same set of experiments, comparing the key subpopulations of both CD8 and CD4_conv_ T cells previously described between ATRi/RT, ATRi/RT/anti-PD-L1 and ATRi/RT/anti-NKG2A/PD-L1 treatment groups. Interestingly, blocking NKG2A was critical in shifting the cell cycle in CD8 T cells from G0/G1 to G2/M phases, and the same effect was seen in CD4_conv_ T cells (figure S4A). These data were further reinforced by the significant increase of Ki67^+^ cells in the treatment cohort that included anti-NKG2A (figure S4B). Adding anti-NKG2A also enhanced the frequencies of both PD-1^+^ CD8 and CD4_conv_ T cells and ICOS^+^ CD4_conv_ T cells (figure S4C-D). Anti-NKG2A antibody significantly promoted the effector memory phenotype, which correlated with increased proportions of Ki67/PRF1^+^ CD8 and CD4_conv_ T_EM_ cells (figure S4E-F). Finally, the inclusion of anti-NKG2A in combination therapy crucially elevated the frequencies of NKG2D- and T-BET-positive CD8 and CD4_conv_ T cells (figure S4G-H).

Taken together, these data indicate that ATRi/RT/anti-NKG2A/PD-L1 combination therapy leads to the infiltration of PD-1^high^/cytotoxic/proliferative/effector memory CD8 and CD4_conv_ T cells in the TME. In parallel and importantly, the nature of CD4_reg_ T cell populations following ATRi/RT is not affected by the addition of anti-NKG2A/PD-L1 immune checkpoint blockade.

### CD40 signalling mediates antitumour immunity in response to ATRi/RT/anti-NKG2A/PD-L1 combination therapy

As described above, prominent subpopulations of both CD8 and CD4_conv_ T cells in the TME following ATRi/RT/ImmuT were characterised by high surface expression of PD-1. We detected in similar experiments enhanced expression of CD40L on those PD-1^+^ T cells i.e., the most activated cells (figure 5A). In parallel, the same treatment conditions led to an increase of CD40^high^ cDC1, cDC2 and pDCs, as well as CD40^+^ monocytes, macrophages and neutrophils (figure S5A-B). These findings prompted evaluation of whether CD40/CD40L signalling was a potential costimulatory pathway responsible for the therapeutic effect seen.

**Figure 5.**
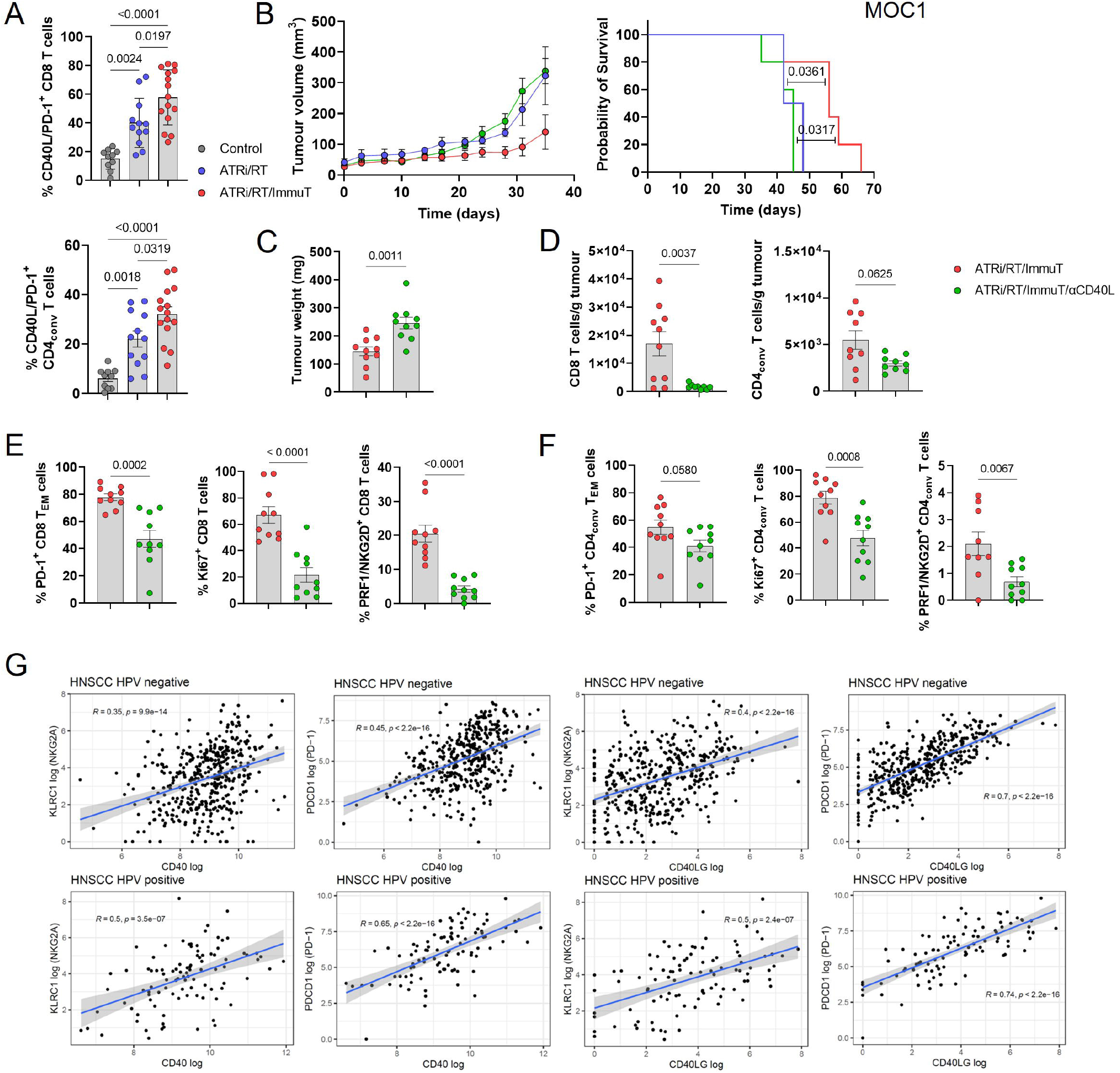
CD40 signalling mediates the efficacy of ATRi/RT and immunotherapy combination. Experiments presented in this figure were performed in the MOC1 model. (A) % of PD-1/CD40L double positive CD8 and CD4_conv_ T cells across indicated treatment conditions (10-15 mice/group). (B) Tumour growth and survival curves across the different conditions (5 mice/group). (C) Tumour weight collected from mice in the indicated conditions (10 mice/group). (D) Absolute number/gram of tumour tissue of CD8 and CD4_conv_ T cells in the various conditions (10 mice/group). % PD-1^+^, Ki67^+^ and PRF/NKG2D^+^ CD8 (E) and CD4_conv_ (F) T cells in treatment conditions (10 mice/group). (G) TCGA database analyses of the correlation between expression of *KLRC1* and *PDCD1* with *CD40* and *CD40LG* in both HPV^-^ and HPV^+^ HNSCC patients.

Indeed, we found that stimulation of CD40 following ATRi/RT significantly reduced tumour burden (figure S5C), whereas interfering with CD40/CD40L signalling abrogated the additive therapeutic effect of immune checkpoint blockade (figure 5B-C). Furthermore, blocking this costimulatory pathway decreased infiltration of both CD8 and CD4_conv_ T cells (figure 5D) and remaining tumour-infiltrating CD8 and CD4_conv_ T cells lost PD-1 expression, effector memory status, proliferative and cytotoxic capacities (figure 5E-F). Finally, to highlight the translational potential of this findings, analysis of the TCGA database revealed that both *CD40* and *CD40LG* expression significantly correlated with *KLRC1* and *PDCD1* expression in HNSCC patients, regardless of their HPV status (figure 5G).

Taken together, these results demonstrate that CD40/CD40L signalling plays a key role in mediating the full efficacy of ATRi/RT/ImmuT by promoting the infiltration and activation of highly activated, proliferating CD8 and CD4_conv_ T cells.

### Combination of ATRi/RT and anti-NKG2A/-PD-L1 immunotherapy significantly affects the dynamics of TCR activity

To study the temporal progression and dynamics of T cell activity in response to combination therapies, we used the recently developed ‘’Timer and Cell Kinetics and Activity’’ (Tocky) system^25, 26^. Tocky is a reporter system based on a fluorescent ‘Timer’ protein, which spontaneously changes its emission from blue to red within 4 hours (figure 6A). The red-form protein is stable, so its maturation allows flow cytometric-based monitoring of rapid temporal changes in expression. By using transgenic mice in which the expression of fluorescent Timer protein is synchronised with the Nr4a3 gene (Nr4a3-Tocky), we can study the induction of TCR activation. Resting T cells do not express the Timer protein (negative signal) but, once T cells recognize cognate antigen and receive TCR signals, Timer transcription is initiated and they become blue^+^ red^-^ (new signal). T cells with persistent TCR signalling sustain Timer transcription over time and accumulate both blue^+^ and red^+^ proteins (persistent signal). When T cells disengage from their antigens, Timer transcription is arrested, and the unstable blue^-^ protein is lost, resulting in T cells becoming blue^-^ red^+^ (arrested signal) (figure 6A).

**Figure 6.**
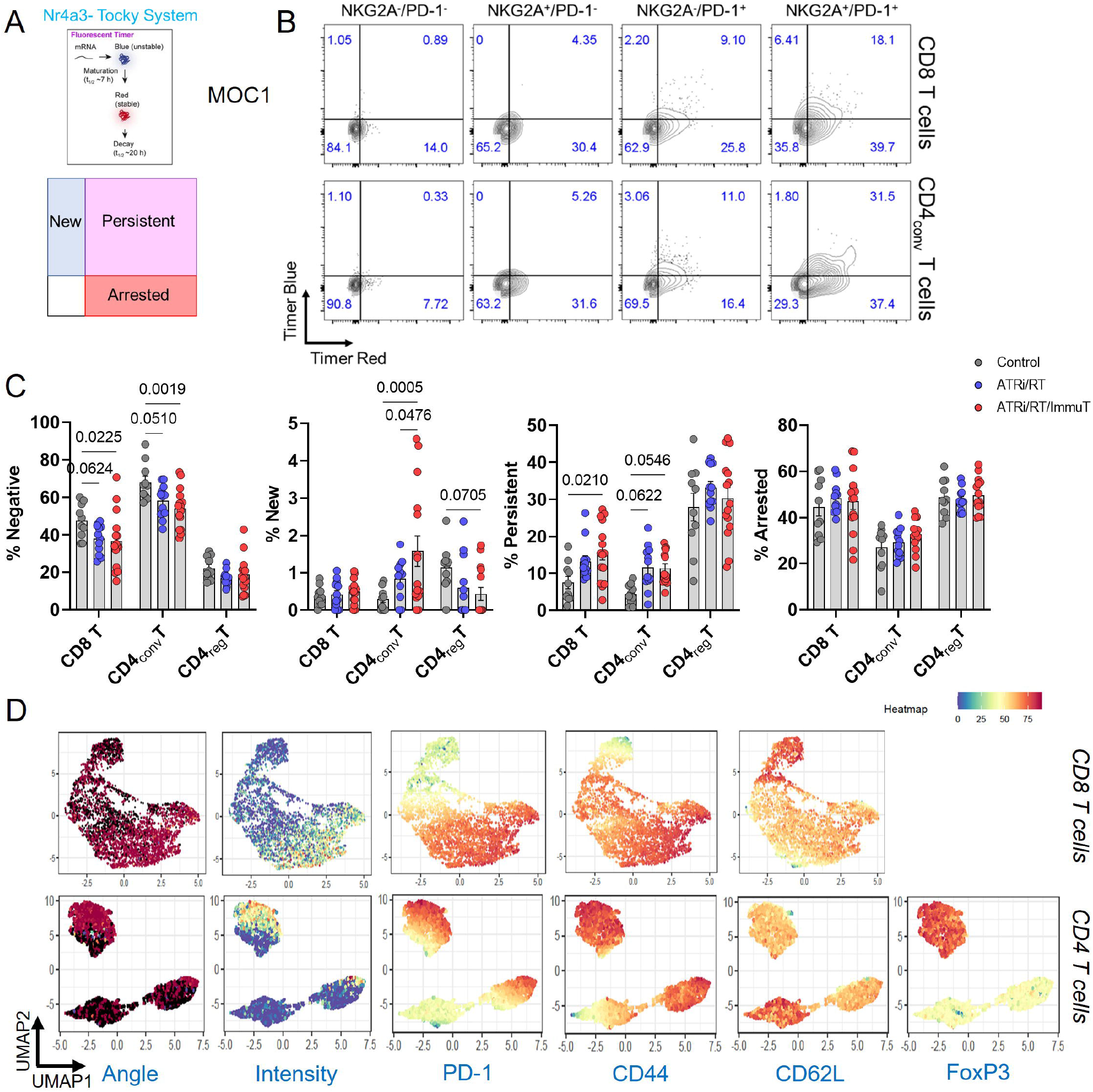
RT, DDRi and anti-NKG2A/PD-L1 immunotherapy modulate the dynamics of TCR activity in both CD8 and CD4 T cells. Experiments presented in this figure were performed in the MOC1 model. (A) Higher panel; schematic explaining production and decay of the Timer protein; lower panel: gating strategy for the detection of the different Timer populations, negative, new, persistent and arrested. (B) Dot plots showing the % of the different Timer population in NKG2A^-^/PD-1^-^, NKG2A^+^/PD-1^-^, NKG2A^-^/PD-1^+^ and NKG2A^+^/PD-1^+^ CD8 and CD4_conv_ T cells (concatenated from 6 mice). (C) % of the indicated Timer population in CD8, CD4_conv_ and CD4_reg_ T cells across all conditions (10-15 mice/group). (D) UMAP plot showing Tocky angle and expression intensity as well as indicated markers expression intensity (concatenated from 14 mice independent of condition).

NKG2A/PD-1 double-positive CD8 and CD4_conv_ T cells were characterised by increased frequencies of persistent TCR activation when compared to other subpopulations (figure 6B). Additionally, CD40L^+^ CD8 and CD4_conv_ T cells were more persistent than the CD40L^-^ population (figure S6A). The action of combined ATRi/RT/ImmuT significantly reduced the proportions of Timer-negative CD8 and CD4_conv_ T cells, which demonstrated enhanced TCR engagement (figure 6C). No variations were observed in terms of new TCR signalling in CD8 T cells across all treatment groups (figure 6C). In contrast, we observed significantly higher proportions of new TCR signals in CD4_conv_ T cells with the addition of ImmuT to ATRi/RT (figure 6C). Whilst there was a trend towards a decrease in the same new Timer population in CD4_reg_ T cells, we detected a significant increase in the frequencies of CD8 T cells with persistent TCR signal in response to ATRi/RT/ImmuT (figure 6C). Trends for increased proportions of persistent CD4_conv_ T cells were found in both ATRi/RT and ATRi/RT/ImmuT treatment groups (figure 6C). Persistence of TCR activation in both CD8 and CD4_conv_ T cells was significantly dependent on CD40/CD40L signalling (figure S6B-C). There was no impact in the frequencies of arrested TCR signalling in all T cells under any treatment condition. Interestingly, populations of CD8 and CD4_conv_ T cells with high Timer intensity were enriched for PD-1 and CD44 but devoid of CD62L, which highlighted their PD-1^+^ effector memory status (figure 6D), as shown by Uniform Manifold Approximation and Projection (UMAP) analysis of Timer Angle and Intensity with markers^27^.

Taken together, these data show unique dynamics of TCR activity from the various populations of T cells in the TME after treatment. Only with the full ATRi/RT/ImmuT combination did CD8 T cells display more persistent TCR activation, while CD4_conv_ T cells exhibited more recent TCR engagement. Importantly, however, the dynamics of TCR activity in CD4_reg_ T cells was not significantly impacted by the different therapeutic regimens, suggesting a more prominent T cell signalling function for CD4_conv_ T cells in cooperation with CD8 T cells.

### Combination of ATRi/RT and anti-NKG2A/PD-L1 immune checkpoint blockade induces variations of TCR repertoires in tumours

To follow up on Tocky analyses and to explore further the amplitude of T cell antitumour response, we subsequently evaluated TCR repertoires in response to different therapy groups. RNA-based targeted sequencing of the CDR3B chain of the TCR was performed from fresh-frozen tumours from mice under three different conditions (control, ATRi/RT and ATRi/RT/ImmuT).

Only the ATRi/RT/ImmuT combination was associated with a significantly higher number of absolute unique productive clonotypes compared to the control, which suggests an increase in TCR repertoire richness (figure 7A). Next, Gini Simpson coefficient was used to quantify clonality across the different conditions. We found that only the full combination showed a significantly higher TCR repertoire clonality compared to the control, suggesting intra-tumoral T-cell expansion (figure 7B). However, no differences were found between the ATRi/RT and the addition of ImmuT. This was also confirmed when exploring the proportion of the repertoire occupied by the first quintile (Q1), which was found to be higher compared to the control in both ATRi/RT and ATRi/RT/ImmuT arms (figure 7C-D). The V- and J-usage did not significantly change across the different conditions, although a trend towards an increase in the even distribution of the V- and J-gene segment pairing was observed when immunotherapy was added to ATRi/RT (figure 7E). When clustering the expanded TCRs by CDR3B triplet amino acid similarity (figure 7F), we found that only the full combination displayed a significant increase in the normalised cluster count compared to the control (figure 7G). Furthermore, these clusters of closely similar TCR clonotypes were not only more abundant, but also bigger in size in the ImmuT-containing combination, as shown when comparing the number of dominant clusters defined as those formed by ≥ 3 expanded clonotypes (figure 7G). This observation strongly supports the notion that the addition of ImmuT to ATRi/RT leads to preferential expansion of large clusters of antigen-sharing T-cell clonotypes, which may reflect enhanced stimulation by increased antigen cross-presentation and T-cell priming, and/or the ability of this combination to promote primed T-cell activation and expansion. Furthermore, the effect of adding anti-CD40L to the triple combination was interrogated. Blocking treatment with anti-CD40L significantly affected the TCR repertoire by abolishing the effect of ATRi/RT/ImmuT in TCR richness (figure S7A), clonality (figure S7B-C) and antigen-sharing clustering (figure S7D-E).

**Figure 7.**
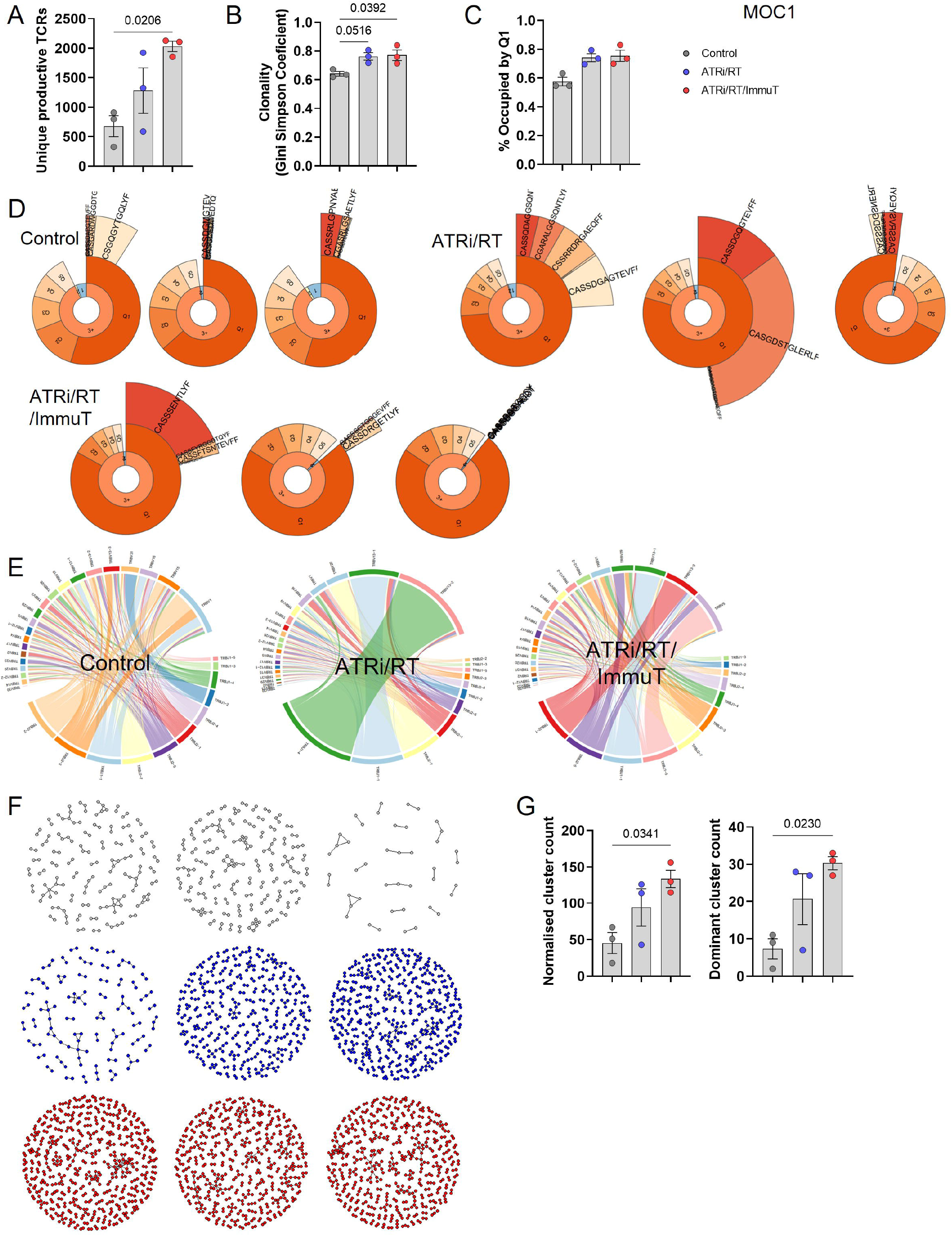
Variations in tumour TCR repertoire following ATR/RT and anti-NKG2A/PD-L1 immunotherapy. (A) Absolute number of unique productive clonotypes across the different conditions. (B) Clonality (Gini Simpson coefficient) across different conditions. (C) Comparison of the proportion (in percentage) of the TCR repertoire occupied by the first quantile of clonotypes. (D) Clonality plots for each condition. The plots present 3 layers to visualize the TCR repertoire clonality: the first layer includes the frequency of singleton (“1”, met once), doubleton (“2”, met twice) and high-order (“3+”, met three or more times) clonotypes; the second layer (“quintile”), displays the abundance of top 20% (“Q1”), next 20% (“Q2”), … (up to “Q5”) clonotypes for clonotypes from “3+” set; and the last layer (“top”) displays the individual abundances of top 5 clonotypes. (E) Representative of V-J junctions by circus plot for each condition. Arcs correspond to different V and J segments, scaled to their frequency in samples and ribbons represent V-J pairings and their size is scaled to their pairing frequency. (F) Network diagrams of CDR3B amino acid triplet clusters for each condition. Clusters containing expanded CDR3s are shown. (G) Comparison of normalized cluster count and dominant cluster count for each condition.

Together, this strongly suggests that the intra-tumoral adaptive immune effects of ATRi/RT and ATRi/RT/ImmuT require engagement of CD40-CD40L signalling, contributing to T cell expansion and TCR repertoire remodeling.

## Discussion

The development of novel therapeutic interventions to eradicate cancer in the future will certainly require, among other approaches, a focus on the most efficient ways to combine targeted therapy, radiation and ICB. Previous clinical trials on the combination of chemoradiotherapy and anti-PD-1/PD-L1 have shown successful results, particularly in the context of lung cancer^28, 29, 30^. However, the outcomes of equivalent strategies in head and neck cancer patients have been disappointing to date^4, 5, 31^ (NCT03040999), probably because they have employed the rather blunt methodological approach of adding anti-PD1/-PD-L1 therapy to standard-of-care (chemo)radiotherapy without any consideration of baseline or on-treatment TME biology. It is highly likely that addition of other immune modulating therapies will be necessary to fine-tune the TME against malignant cells during or following chemoradiotherapy. One such candidate is the anti-NKG2A antibody, monalizumab, which is being investigated in clinical settings^10, 12^. In the present study, we describe the efficacy of a combination of ATR inhibitor, radiation and anti-NKG2A/anti-PD-L1 immune checkpoint blockade in mounting a protective antitumour immune response in both HPV^-^ and HPV^+^ preclinical models of head and neck cancer. Enhanced antitumour functions of NKG2A^+^ CD8 T cells have been described in previous studies. NKG2A was reported to be a late immune checkpoint and NKG2A expression in CD8 T cells was associated with repeated stimulation and cell division^32, 33^. NKG2A-positive clusters of CD8 T cells in head and neck cancer patients were enriched for other immune checkpoints, including PD-1, TIM-3 and CD39^33^. Furthermore, NKG2A^+^ CD8 T cells were characterised by a proliferative, cytotoxic and tissue-resident memory phenotype with TCR-independent functions in bladder cancer^32^. Herein, we extended those findings by showing the presence of highly activated NKG2A/PD-1 double-positive CD8 T cells in human head and neck cancer tumours. Very low numbers of double-positive CD4 T cells were detected. However, analyses in samples from the PATRIOT clinical trial (NCT02223923) showed that ATRi/RT increased the frequencies of both NKG2A/PD-1 double-positive CD8 and CD4 T cells in partial and complete responders. Interestingly, our preclinical murine results were concordant with human scRNAseq with very low NKG2A^+^/PD-1^+^ CD4 T cell numbers in the TME of both non-treated HPV^-^ and HPV^+^ tumours. Mouse studies also allowed us to confirm that ATRi/RT led to an increase in the proportion of both NKG2A^+^/PD-1^+^ CD8 and CD4 T cells in the TME. These findings reinforced the hypothesis that targeting NKG2A/PD-1 double-positive T cells after ATRi/RT could represent a promising therapeutic strategy. Moreover, the development of bispecific blocking antibodies targeting NKG2A and PD-1/PD-L1 axes is likely to facilitate the design of future clinical trials.

In the data reported here, not only have we demonstrated the benefit of ATRi/RT/anti-NKG2A/-PD-L1 on tumour growth and survival in both ectopic and orthotopic models, but we have also elaborated, in detail, the ability of this combination to reconfigure the biology of both CD8 and CD4 T cells in the TME. Dual anti-NKG2A/-PD-L1 immunotherapy, added as an adjuvant to ATRi/RT, drives significant infiltration of PD-1^high^/proliferative/effector memory cytotoxic CD8 and CD4_conv_ T cells in tumours. We found that addition of anti-NKG2A antibody to the combination was essential to shifting the cell cycle towards the G2/M phase in T cells, which was in line with previous data showing repeated cell division in NKG2A^+^ CD8 T cells^33^. These results are also consistent with findings of elevated proliferative capacity of CD8 T cells in response to combined RT and dual NKG2A/PD-1 blockade^34^.

Maximal efficacy of the full combination therapy relied on recruitment and activation of both CD8 and CD4_conv_ T cells. While most previous publications examining NKG2A-targeted immunotherapy in pre-clinical cancer models have been focused on NK and CD8 T cells^7, 8, 35, 36^, to our knowledge, this is the first description of an anti-NKG2A blocking antibody interacting with both CD8 and CD4_conv_ T cells to optimise its therapeutic benefits. Previous investigations identified a critical role of CD4_conv_ T cells in helping CD8 T cells to achieve tumour rejection^37^. Indeed, it was reported that IL-2-producing CD4 helper T cells increased the number and cytolytic function of tumour-specific CD8 T cells^38^. Arina and colleagues also showed that adoptive transfer of alloantigen-specific CD4 T cells rescued exhausted neoantigen-specific CD8 T cells, leading to eradication of relapsing tumours^39^. To increase our understanding of the mechanisms behind cooperation between CD8 and CD4 T cells, the antibody-mediated depletion and Tocky timer system data provided insights into the dynamic contribution of T cells, including CD4_conv_, in the TME, to the overall therapeutic effect following ATRi/RT/ImmuT combination. The impact of the double depletion of CD8/CD4 T cells on tumour growth, when compared with the depletion of CD8 T cells alone, correlates well with the data on accumulation of CD4_conv_ T cells with new TCR activation in the tumour bed. Together, these data support the view that newly activated CD4_conv_ T cells sustain effective antitumour immune responses in the TME by supporting more persistent CD8 T cell activity.

Our restimulation assay, using an HPV long-overlapping peptide, provides compelling evidence of an augmented level of antigen specificity that develops in mice following ATRi/RT and combination immune checkpoint blockade. An additional contributor to the effectiveness of our therapeutic approach could be its favourable impact on the intra-tumoral TCR repertoire. While the ATRi/RT combination alone exhibits a profound influence on the expansion of T cell clonotypes within the TME, our observations reveal a notable augmentation in the richness of the TCR repertoire and the emergence of numerous and large antigen-sharing TCR clusters in response to the ATRi/RT/ImmuT regimen. These findings collectively imply that the full combination is essential to achieving an optimized immune response against tumours due to its concomitant enhancement of TCR repertoire richness, clonality, and antigen specificity. In a pre-clinical breast cancer study, Rudqvist et al. reported that RT broadened the TCR repertoire and that combination of RT with anti-CTLA-4 antibody triggered T cell clone expansion and increased CDR3 clustering compared to control^40^. Moreover, differential response to anti-PD-L1 treatment has recently been attributed to variations in the TCR repertoire in a preclinical model of HNSCC^41^. Previously published human studies around TCR repertoire changes, secondary to immunotherapy or combination of immunotherapy and RT, have shown that increased TCR repertoire clonality and clustering are both correlated with response to immunotherapy and, therefore, desirable outcomes of combination therapeutic regimens^42, 43, 44^. In addition, Wand et al. documented heightened TCR diversity in the peripheral blood of responders among patients with HNSCC who underwent combined treatment involving cetuximab and nivolumab^45^.

The ability of CD40 stimulation to support T cell antitumour responses has been extensively documented^46^. However, clinical development of CD40 agonist therapies have, so far, been significantly hindered by the occurrence of dose-limiting toxicities^47^. Our findings suggest that dual NKG2A/PD-L1 blocking intervention following ATRi/RT could mimic the efficacy of CD40 agonist therapy without recapitulating its significant and limiting toxicity. To add credibility to this hypothesis, our results show that anti-NKG2A/PD-L1 combination treatment following ATRi/RT increases the expression of CD40 on myeloid cells as well as CD40L on T cells, these two events subsequently mediating the recruitment, activation and proliferation of CD8 and CD4_conv_ T cells in the TME. We show here that CD40/CD40L signalling is important in sustaining persistent TCR activation in CD8 T cells in response to ATRi/RT and immunotherapy combination. Meanwhile, our TCRseq analyses are in line with previous work describing an important function of this signalling pathway in affecting the TCR repertoire^48^.

To conclude, in the present study, we unveil a means to redirect the full arsenal of T cell responses towards optimal cancer cell eradication in both HPV^-^ and HPV^+^ tumours. This work provides a clear rationale for designing future clinical trials of radiation, DDRi and dual immune checkpoint blockade-based immunotherapy for patients with head and neck cancer.

## Methods

### Human studies

Two single-cell RNA sequencing datasets from recently published studies^19, 20^ were analysed to explore immune cell types expressing *KLRC1/PDCD1* in head and neck squamous cell carcinomas (HNSCC). Both datasets include single-cell whole transcriptomes from HPV+ and HPV-HNSCC tumours and from peripheral blood; Cillo A et al. data also includes CD45^+^ cells sorted from 5 tonsil tissues from independent patients without cancer. Gene/barcode matrices for each patient generated by CellRanger pipeline were downloaded from GSE164690 and GSE181919. Each dataset was processed separately in R using ‘Seurat’ package (v4.3.0). Cell-based and gene-based quality control filters were applied. Firstly, cells with less than 1000 transcripts (UMIs) or with less than 200 genes per cell were filtered out from further analysis. Next, cells that expressed more than 5000 genes or with ratio of mitochondrial transcripts higher than 10% per cell were also excluded. Finally, only genes expressed in at least two cells in each dataset were kept. Both datasets were than normalized using ‘SCTransform’ (with mitochondrial transcripts ratio effect regressed out), followed by dimensionality reduction and Louvain clustering. Cell-type cluster annotation was performed using sets of marker genes: CD3D with combination with CD4, CD8A and FOXP3 for CD4^+^, CD8^+^ and regulatory T cells; KLRD1 and KIR2 for NK cells; CD19 and CD79A for B / plasma cells; CD14, CD68, FCGR3A and IL3RA for myeloid populations; TPSB2 and TPSAB1 for mast cells; EPCAM, KRT14 and KRT17 for epithelial cells; PECAM1 and RAMP2 for endothelial cells; COL1A1, COL1A2 and DCN for fibroblasts; RGS5 for pericytes. For the TCGA analyses, we downloaded clinical data and RSEM (RNAseq by Expectation-Maximamization) normalised expression data for the Head and Neck squamous cell carcinoma (HNSC) cohort of TCGA from the Firebrowse website [http://firebrowse.org/] version 2016012800.0.0. We generated the graphs using R package “ggplot”. Statistic is Spearman R. We used R version 4.3.1. Blood samples were taken from patients enrolled on the phase I PATRIOT study of Ceralasertib (AZD6738) in combination with palliative radiotherapy (EudraCT: 2013-003994-84). Patients were 18 years and over, with a diagnosis of advanced solid malignancy, and an indication for palliative radiotherapy. All patients were ECOG performance status 0-2 with a life expectancy of at least 3 months and adequate organ function. This study was conducted in accordance with protocol requirements, Good Clinical Practice, the guiding principles of the Declaration of Helsinki All participants provided written informed consent. The protocol was reviewed by the local ethics committee. Radiotherapy was administered daily, 2 Gy per fraction, Monday-Friday. A total of 30 Gy in 15 fractions was administered. Ceralasertib was administered orally at a dose of 80 mg (the recommended phase II dose), twice daily, for 3-7 days before the start of radiotherapy, during radiotherapy, and for 2 days after. Blood was taken before Ceralasertib dosing, before fraction 1, before fraction 6, and before fraction 11, in an 8 mL EDTA tube and was processed within 24 hours of sampling for flow cytometry.

### Mouse models of head and neck cancer

All animals were handled according to the Institute of Cancer Research and U.K. Home Office guidelines. In this study, we used models of murine immunocompetent HPV-negative and HPV-positive head and neck cancer, MOC1 and mEER, respectively. The C57BL/6 mouse cancer cell line MOC1 and mEER was kindly provided by Ravindra Uppaluri (Dana-Farber Cancer Institute) and Paola Vermeer (Sanford Research), respectively. Cells were cultured and maintained in Dulbecco Modified Eagle Serum (DMEM) supplemented with 10% foetal calf serum (FCS), 2mM L-glutamine, penicillin/streptomycin and sodium pyruvate. 8- to 14-week-old females C57BL/6 WT (purchased from Charles River) or Nr4a3-Tocky: Foxp3-EGFP mice^26, 49^ were inoculated subcutaneously (ectopic) on the right flank with 4 × 10^6^ cells for the MOC1 or 10^6^ cells for the mEER model. In some experiments, mice were inoculated in the lip / cheek at the entrance of the oral cavity (orthotopic) with 3 × 10^6^ MOC1 cells. Tumour growth was monitored every 3/4 days using a calliper and expressed as volume ((length × width × depth)/2). Mice were euthanised when tumours reached an average of 15 or 12 mm in two of the three dimensions for the ectopic or orthotopic models, respectively. When indicated, cured animals were rechallenge subcutaneously 90 days after the first challenge.

### In vivo tumour treatments

*In vivo* treatments were initiated when tumours reached an average of 20-100 mm^3^. The ATR inhibitor AZD6738 was provided by AstraZeneca and solubilised in 10% DMSO, 40% propylene glycol, 50% water, orally administrated 2 hours before irradiation for 4 consecutive days at 75 mg/kg/dose as indicated. Animal tumours were irradiated under anaesthesia with ketamine/xylazine injected intraperitoneally. Irradiation was initiated one day after ATR inhibitor dosing and was performed using an AGO 250kV X-ray machine at a dose rate of 1.62 Gy/minute (AGO). Animals were irradiated in the prone position positioned underlead shielding with a 16-mm diameter aperture aligned over the Tumour. A total of 9 (RNAseq, flow cytometry and TCRseq experiments) to 18 Gy (survival experiments) radiation dose in three fractions were given. Radiation dose was measured using a Farmer Chamberand Unidos-E Dosemeter (both PTW). Three days after the last dose of radiotherapy, mice were treated with anti-NKG2A (200 μg/mouse i.v., every 4/5 days for two weeks for a total of three injections) and/or anti-PD-L1 antibodies (200 μg/mouse i.p., every ¾ days for two weeks for a total of 4 injections) provided by AstraZeneca. For the depletion of NK cells, mice were injected intraperitoneally with 200 µg/mouse of anti-NK1.1 antibody (BioXcell, clone PK136), twice weekly starting 1 day before treatment. For the depletion of CD8 T cells, mice were injected intraperitoneally with 200 µg/mouse of anti-CD8α (BioXcell, clone 2.43) twice weekly starting 1 day before treatment. For the depletion of CD4 T cells, mice were injected intraperitoneally with 500 µg/mouse of anti-CD4 (BioXcell or Biolegend, clone GK1.5) for the first injection starting 1 day before treatment and 250 µg/mouse twice weekly thereafter. For the analysis of cell cycle position, mice were injected i.p. with 1 mg of BrdU (BD Biosciences) 12 hours before tumour harvest.

### RNA sequencing

MOC1 and mEER tumours were harvested 10 days after the last radiation dose. RNA from tumours was extracted and purified from tumours using the NucleoSpin RNA extraction kit (Macherey-Nagel). Library preparation and sequencing were performed by Genewiz (Leipzig, Germany). Library prep was strand-specific, used PolyA selection, with 2x150bp sequencing at 10 million pairs performed on an Illumina NovaSeq. Quality control was performed using FastQC. Low quality bases and adaptor sequences were removed using Trimmomatic. Reads were mapped to the M. musculus GRCm38 reference genome using HISAT2, samtools and StringTie. Differential gene expression was assessed using DESeq2. Active-subnetwork-orientated gene set enrichment analysis using STRING protein-protein interaction networks and Kyoto Encyclopedia of Genes and Genomes (KEGG) pathways was performed using pathfindR. Immune cell composition was estimated from transcriptomic data using mMCP-counter. Data visualization utilized ggplot2 and ComplexHeatmap packages.

### Flow cytometry analysis

Harvested tumours were excised, finely minced, and enzymatically digested for 30-45 min at 37°C in PBS containing 1 mg/ml collagenase type VI (Sigma-Aldrich) 1 mg/ml DNase type I (Roche), 100 ng/ml Dispase (Sigma-Aldrich) and Trypsin. Digested tumours were smashed and filtered through 70 μm-pore cell strainers. Cells were centrifuged at 1400 rpm at 4°C for 10 min and pellets washed in PBS/2% FCS/5mM EDTA. Harvested spleen were mechanically dissociated, smashed and filtered through 70 μm-pore cell strainers. Red blood cells were lysed by a 1 min incubation in ACK lysis buffer (Thermofisher Scientific). Cells were centrifuged at 1400 rpm at 4°C for 10 min and pellets washed in PBS/2% FCS. For some experiments, mice were bled 20 µL from the tail vein. PBMCs from human blood samples were isolated on a density gradient medium (Lymphoprep, STEMCELL Technologies). Cells were incubated 4°C for 10 min with anti-mouse or anti-human CD16/CD32 (BD Pharmingen) prior to surface staining, 4°C for 30 min. Dead cells were excluded using the Fixable Viability Dye eFluor™ 780 (Thermofisher Scientific). The complete list of antibodies used in this study can be found in the supplementary information. Intranuclear detection of Ki67, FoxP3 and T-BET was performed using The Foxp3 Transcription Factor Staining Buffer Set (Thermofisher Scientific) following the manufacturer’s instructions. For surface detection of CD107a, cells were incubated in complete IMDM containing Golgi Plug/Golgi Stop (BD Biosciences) in the presence of a stimulation cocktail containing PMA and Ionomycin (Invitrogen) and mouse CD107a antibody for 4 h at 37°C. Following surface antibody staining, cells were fixed using IC Fixation Buffer (Invitrogen), permeabilized in Permeabilization Buffer (Invitrogen), and stained with mouse IFN-γ antibody. Cell cycle position analyses were performed using the BD PharmigenTM BrDU Flow Kits (BD Biosciences) following the manufacturer’s instructions. Samples were analysed on a LSR II or FACSymphony A5 (BD Biosciences) and FACS analyses were performed using the FlowJo version 10 software. Tocky timer data was analysed using the “TockyAnalysis” package in R (v 3.6.3).

### TCR sequencing

TCR β-chain sequencing was performed by utilizing whole RNA extracted from tumor samples by using a quantitative experimental and computational TCR sequencing pipeline described previously [Oakes et al. Front Immunol, 2017]. Briefly, the TCR library preparation protocol consists of first a T4 ligation step to the 3’ end of the TCR cDNA which allows the amplification of all possible rearrangements using a single set of primers per locus and the incorporation of a UMI attached to each cDNA TCR molecule that enables correction for PCR and sequencing errors. The final library was prepared for sequencing according to the Illumina NovaSeq protocol immediately prior to loading. Sample was loaded at 1.2 pM with 15–20% of PhiX DNA at 1.8 pM. All samples were run in parallel, yielding 5 million reads per sample (2x150 bp paired end). The bioinformatic pipeline used for TCR identification, error correction and CDR3 extraction was Decombinator (freely available at https://github.com/innate2adaptive/Decombinator). For TCR analysis, clonality was calculated using the Gini coefficient.

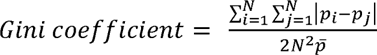

where *p_i_* and *p_j_* represents the frequency of the respective i^th^ and j^th^ sequences in the repertoire and *p* the average of the clone frequencies. Gini coefficient ranges from zero (maximal diversity of the repertoire or equal abundance of each sequence) to one, with high value representing extreme inequality (i.e. high clonality towards one sequence). Therefore, the Gini coefficient increases as the number of abundant clones rises, thus further reducing the frequency of less represented clones (i.e. higher inequality). Moreover, with increasing richness of repertoires, the inequality between dominant and subdominant (low frequency) clones gets wider, leading to a small rise in the Gini coefficient. This coefficient exhibits relatively lower sensitivity to undersample due to the exclusion of unique TCR sequences present in the total repertoire and consequently helps mitigate the inequality introduced by changes in frequency distribution. Unique productive clonotypes and clonality were compared across groups using One-way ANOVA with Dunnett’s multiple comparisons tests. VJ genes junction circles and sample clonality plots were done using the VDJtools bioinformatic software^50^. Proportion occupied by quantile 1 across groups was compared using One-way ANOVA with Dunnett’s multiple comparisons tests. For TCR clustering, the pairwise similarity between pairs of TCRs was measured on the basis of amino acid triplet sharing. Sharing was quantified using the normalized string kernel function stringdot (with parameters stringdot (type = ‘spectrum’, length = 3, normalized = TRUE) from the Kernlab package. The kernel is calculated as the number of amino acid triplets (sets of three consecutive amino acids) shared by two CDR3s, normalized by the number of triplets in each CDR3 being compared. The TCR similarity matrix was converted into a network diagram by using the iGraph package in R. Two TCRs were considered connected if the similarity index was >0.7. Per sample, we counted the number of clusters containing an expanded CDR3. An expanded CDR3 was considered if it was present above a threshold frequency of 2/1,000 (corresponding to the top 1% of the empirical TCR frequency distribution). At this threshold, which we already described in previously published work [Barennes et al., 2020], the correlation between clonality and proportion of repertoire occupied by expanded TCRs is very strong and the number of TCRs labelled as expanded is greater than for higher thresholds for which this correlation is also significant, which enables to keep the greatest amount of data whilst still applying a stringent filtering step. To normalize the counts of clusters obtained for the input size, for each sample, we randomly selected, outside of the real clustering structure, the number of CDR3s equal to the number of expanded CDR3s in that sample and looked for clusters around those. A cluster was considered to be dominant if it was formed by more than 3 CDR3s.

### Statistical methods

Statistical analyses were performed using GraphPad Prism software version 8 and 9. Data are presented as means ± SEM and are from a single or cumulative of two to three independent experiments. Area under curves were used to determine statistical differences in tumor growth between groups. Statistical differences between survival groups were determined using the Log-rank Mantel Cox test. Intensity of marker expression was shown as geometric fluorescence mean intensity (MFI) in the positive population. Unpaired T-test or one-way ANOVA followed by Tuckey’s post T-test were used for statistical analysis when two or multiple groups were analyzed, respectively. When data did not follow a Gaussian distribution, we used nonparametric Mann–Whitney *U* or Kruskal-Wallis tests followed by Dunn’s post T-test for statistical analysis when two or multiple groups were analyzed, respectively. We used two-way ANOVA followed by Tuckey’s multiple comparison test when two parameters were studied. Significant outliers were removed using the Grubb’s test.

