## Supplementary material for "Sculpting the tumour microenvironment by combining radiotherapy and ATR inhibition for curative-intent anti-PD-L1- and anti-NKG2A-based adjuvant immunotherapy"

**Supplementary figures**

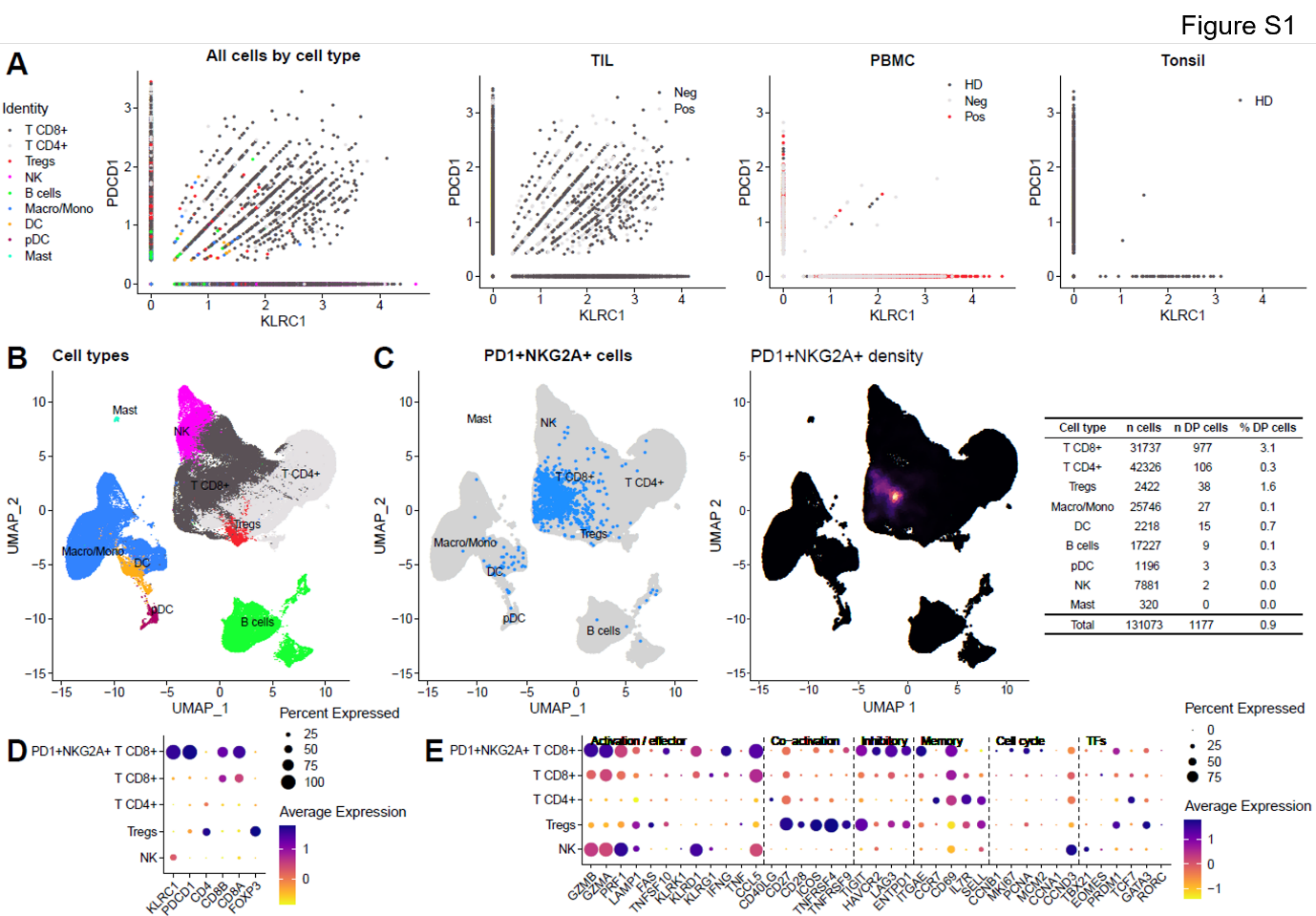

**Figure S1. Highly activated NKG2A and PD-1 double-positive T cells are detected in head and neck cancer patients.** (A) Scatter plots showing expression of *KLRC1* and *PDCD1* in, left panel; various cell population and other panels; tumours (TIL) versus blood (PBMC) and tonsil (B) UMAP plot showing distribution of identified cell clusters distribution. (C) Left panel; UMAP plot depicting NKG2A/PD-1 double positive cells in the different identified cell clusters, right panel; UMAP plot depicting intensity expression of NKG2A/PD-1 double positive (DP) cells with absolute numbers for each identified cell populations (right table). Dot plots showing average and percentage expression of the *KLRC1*, *PDCD1* and pan-cell markers (D) and activation/effector, co-activation, inhibitory, memory, cell cycle and transcription factors (TFs) markers (E) in NKG2A^+^/PD-1^+^ CD8^+^ T cells versus CD8^+^T, CD4^+^T, Tregs and NK cells. Datasets from Cillo A et al., 2020.

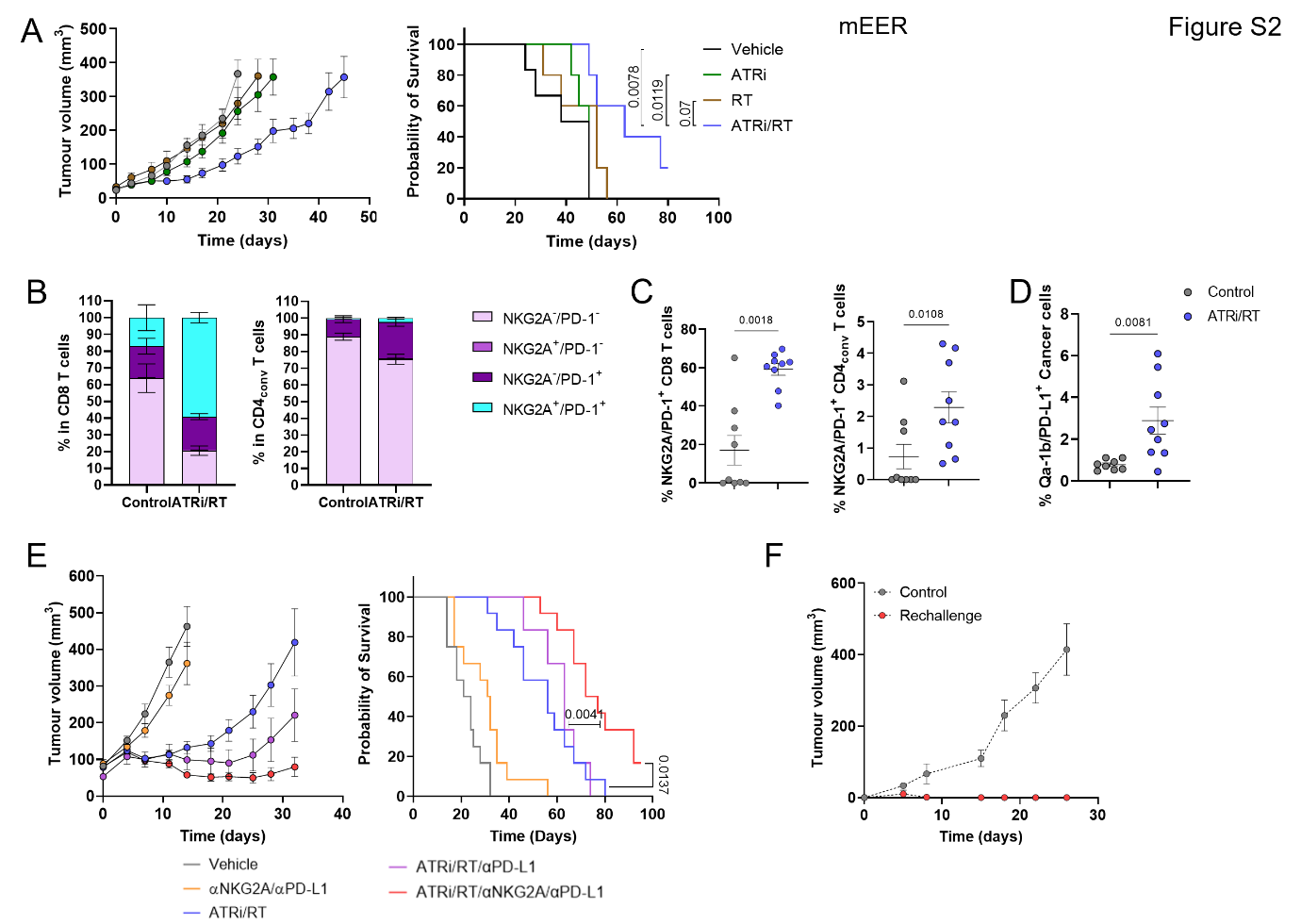

**Figure S2. NKG2A and PD-L1 dual immune checkpoint blockade improves the therapeutic outcome of radiation and DDRi.** Experiments in this figure were performed in the mEER model. (A) Tumour growth and survival curves across the different conditions (5-6 mice/group). (B) % of NKG2A and/or PD-1 positive populations in CD8 and CD4_conv_ T cells (9 mice/group). (C) % of NKG2A/PD-1 double positive populations in CD8 and CD4_conv_ T cells (9 mice/group). (D) % surface expression of Qa-1b/PD-L1 double positive cancer cells in the different conditions. (E) Tumour growth and survival curves across the different conditions in (6-12 mice/group). (I) Tumour growth in control versus rechallenged mice (3-6 mice/group).

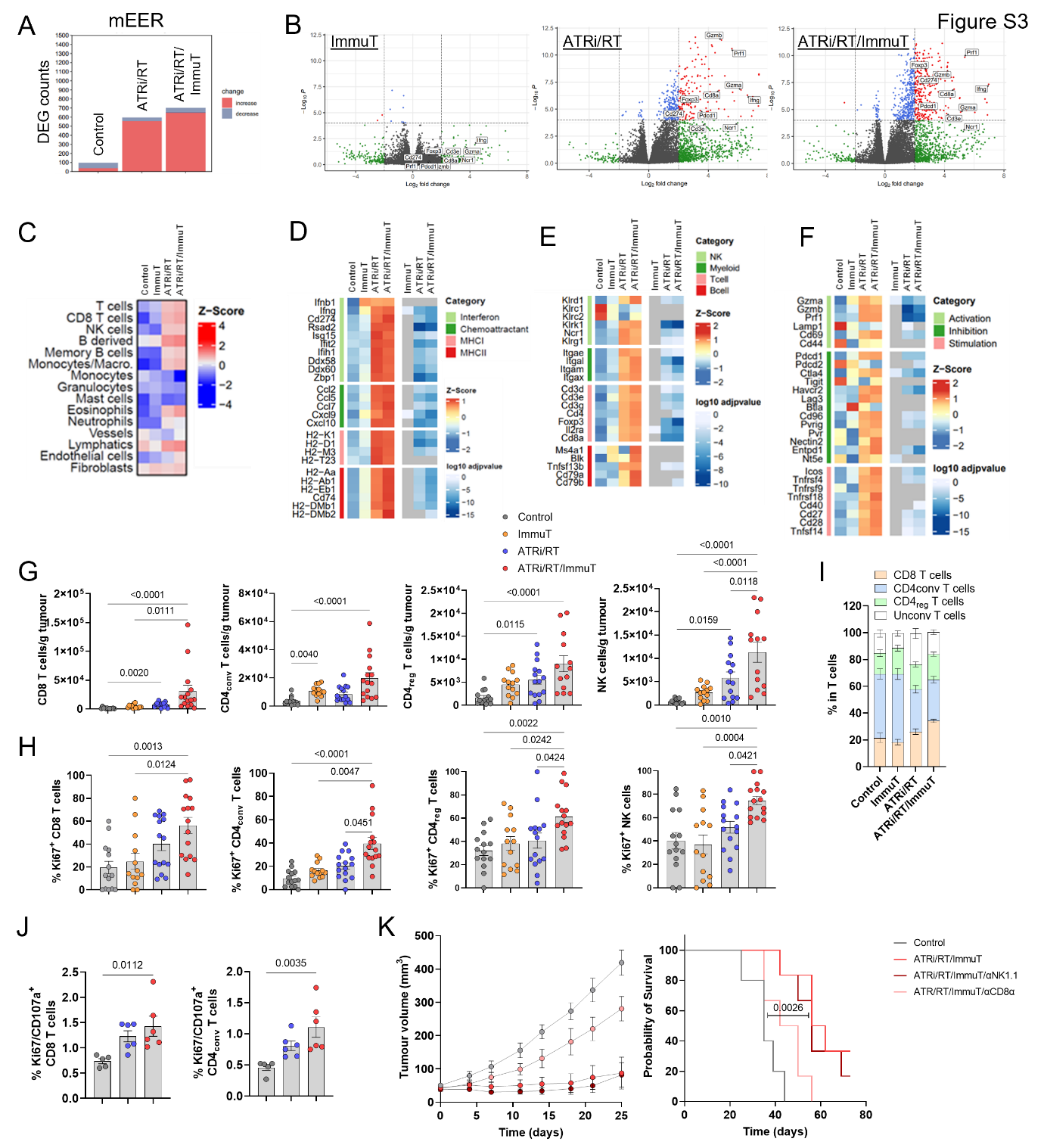

**Figure S3. ATRi/RT and dual anti-NKG2A/PD-L1 immunotherapy trigger a potent CD8 T-cell antitumour immune response in an HPV-positive head and neck cancer model.** Experiments presented in this figure were performed in the mEER model. (A) Number of differentially expressed genes (DEGs) across the different conditions calculated by DESeq2. (B) Volcano plots showing expression of the different genes in the various conditions (genes characteristic of an immune response are named). (C) Immune cell population estimates. Immune cell scoring was performed on normalised RNAseq counts using the mMCP-counter package. Heatmaps corresponding to interferon and cytokine signalling, chemoattractant, MHCI and MHCII (D); immune cell populations (E); and immune cell activation status (F). Data shown are z-scores of log2 transformed normalised counts for the treatment conditions shown. This plotted alongside log10 adjusted *p*-value for each gene calculated from DEG analysis using DESeq2. Non-significant adjusted *p*-values > 0.05 are indicated as grey. (G) Absolute number/gram of tumour of the indicated lymphocytes in the various conditions (13-15 mice/group). (H) % Ki67 positive cells in the indicated lymphocytes in the different conditions (13-15 mice group). (I) % of indicated lymphocytes in the total T cell population across all treatment conditions. (J) % Ki67/CD107a double positive CD8 and CD4_conv_ T cells in splenocytes from mice treated with the indicated conditions restimulated with an HPV peptide (5-6 mice/group). (K) Tumour growth and survival curves across the different conditions (5-6 mice/group).

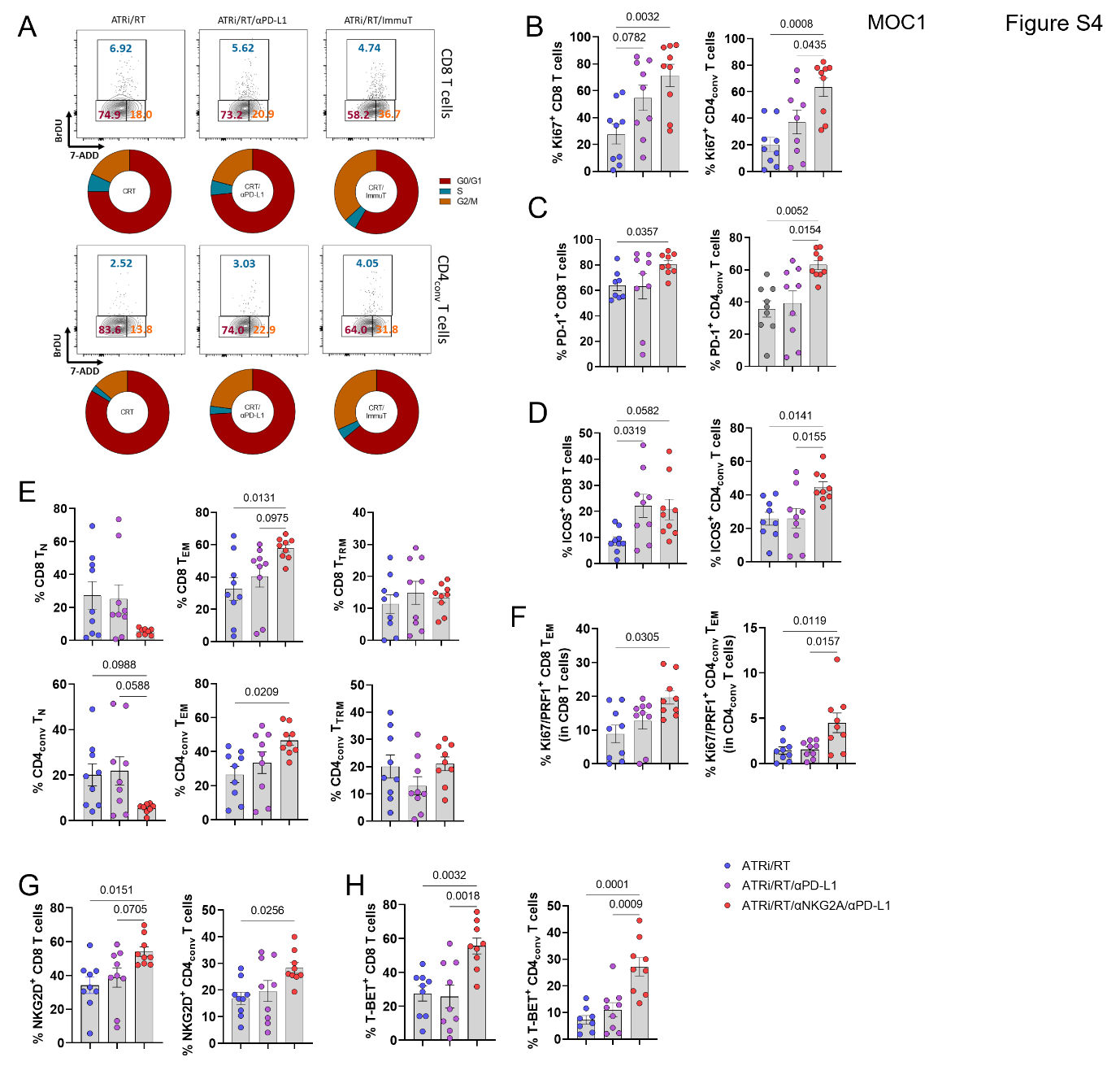

**Figure S4. ATRi/RT and dual anti-NKG2A/PD-L1 immunotherapy induce the proliferation of PD-1^+^ effector memory cytotoxic CD8 and CD4 T-cells in the tumour microenvironment.** Experiments presented in this figure were performed in the mEER model. (C) Dot plot and donut chart representing % of CD8 and CD4_conv_ T cells in the different cell cycle phases across all conditions (concatenated from 6 mice/group). Across all conditions the following graphs show % (B) Ki67^+^, (C) PD-1^+^, (D) ICOS^+^, (E) naïve (N), effector (EM) and tissue-resident (TRM) memory, (F) Ki67/PRF^+^ EM, (G) NKG2D^+^ and (H) T-BET^+^ CD8 and CD4_conv_ T cells (15-17 mice group).

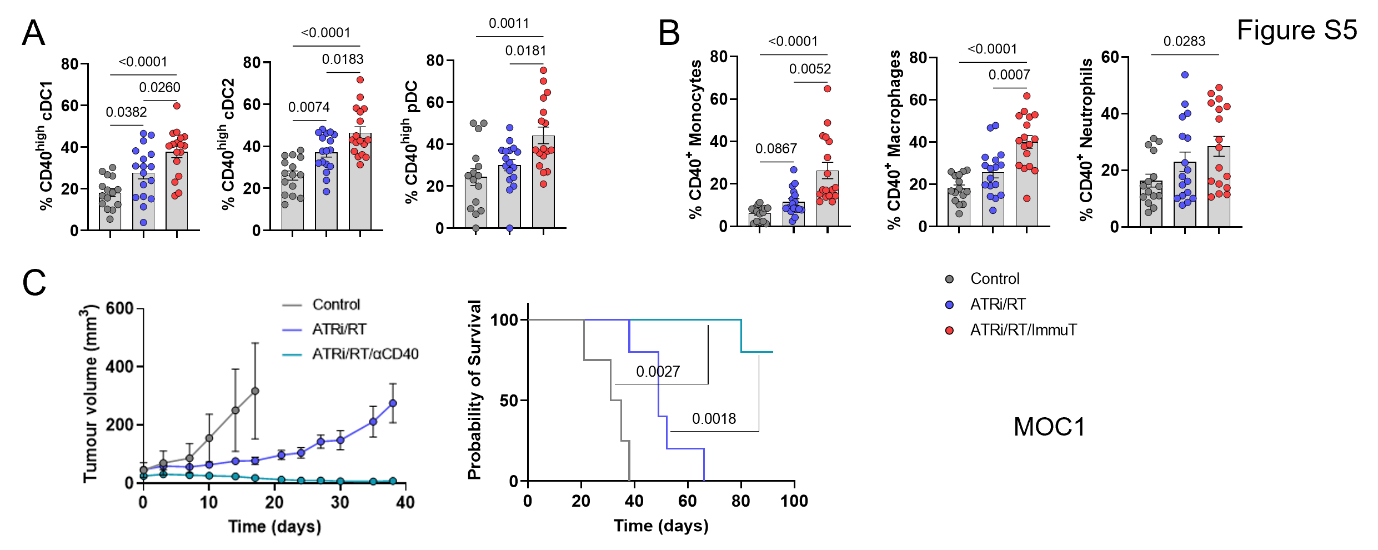

**Figure S5. CD40 stimulation promotes potent antitumour response following ATRi/RT.** Experiments presented in this figure were performed in the mEER model. (A) % of CD40^high^ cDC1, cDC2 and pDC and (B) CD40^+^ monocytes, macrophages and neutrophils across all treatment groups (15-17 mice/group). (C) Tumour growth and survival curves across the different conditions (4-6 mice/group).

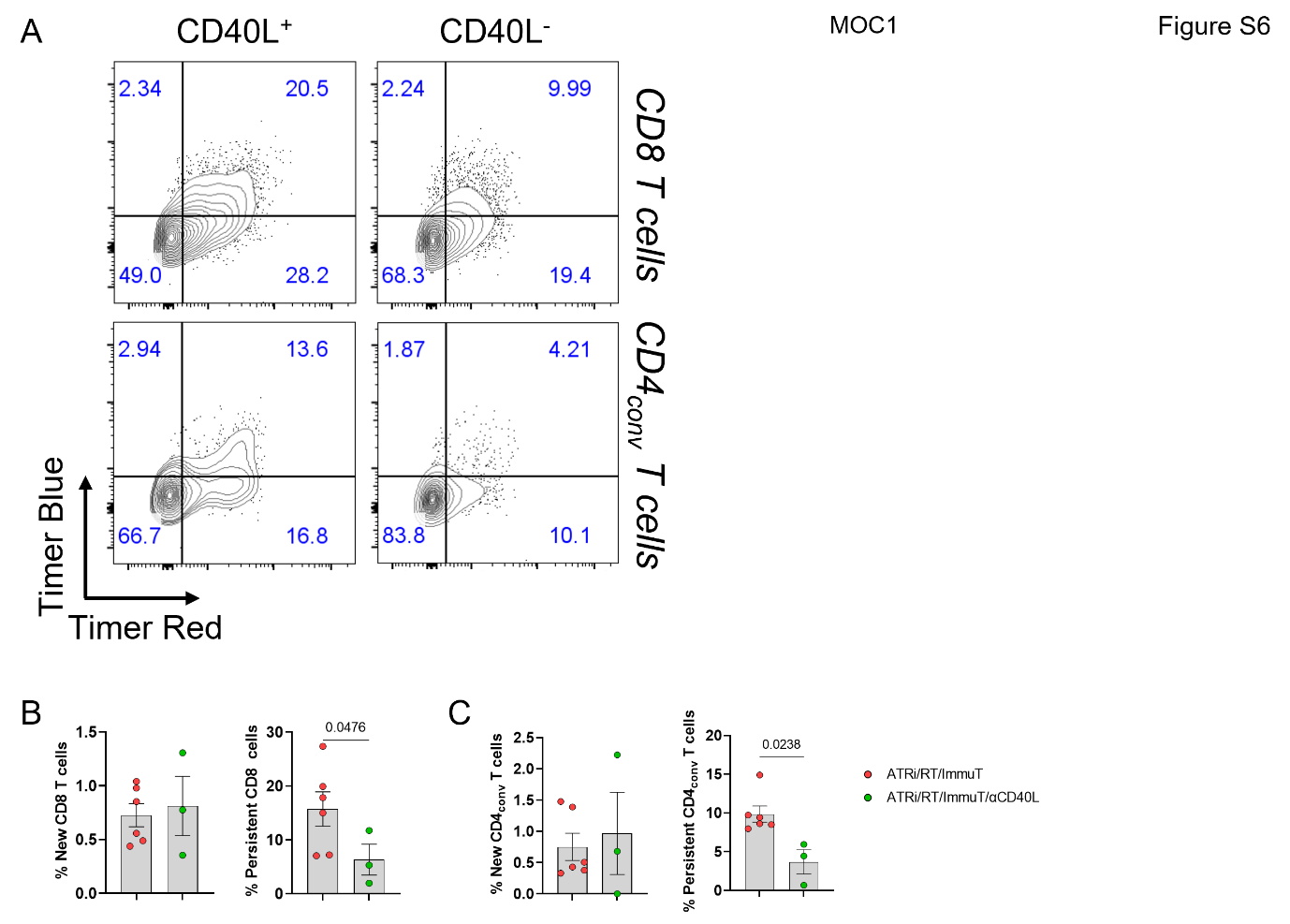

**Figure S6. CD40/CD40L axis mediates the persistence of TCR signalling in both CD8 and CD4_conv_ T cells following ATRi/RT and anti-NKG2A/PD-L1 immunotherapy.** Experiments presented in this figure were performed in the MOC1 model. (A) Dot plots showing the % of the different Timer population in CD40L^+^ versus CD40L^-^ CD8 and CD4_conv_ T cells (concatenated from 6 mice). % of the indicated Timer population in CD8 (B) and CD4_conv_ (C) T cells in both conditions (3-6 mice/group).

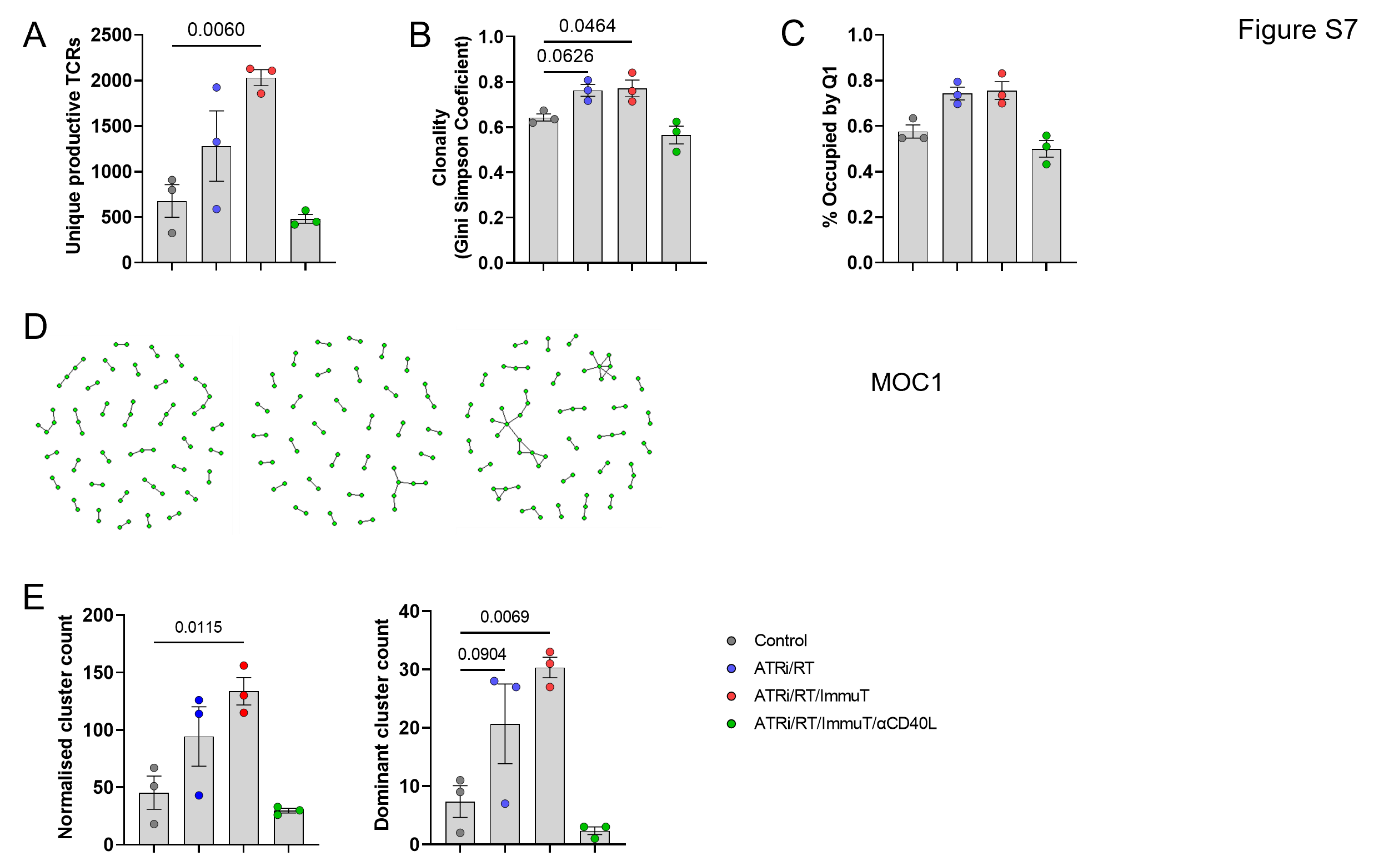

**Figure S7. CD40/CD40L axis mediates the variations in tumour TCR repertoire following ATR/RT and anti-NKG2A/PD-L1 immunotherapy. (**A) Absolute number of unique productive clonotypes across the different conditions. (B) Clonality (Gini Simpson coefficient) across different conditions. (C) Comparison of the proportion (in percentage) of the TCR repertoire occupied by the first quintile of clonotypes. (D) Network diagrams of CDR3B amino acid triplet clusters for ATRi/RT/ImmuT/αCD40L condition. Clusters containing expanded CDR3s are shown. (E) Comparison of normalized cluster count and dominant cluster count for each condition.

**Supplementary table**

Table S1. List of flow cytometry antibodies.

| Epitope | Clone | Fluorophore | Manufacturer |
| --- | --- | --- | --- |
| Anti-human CD45 | HI30 | BV650 | Biolegend |
| Anti-human CD3 | HIT3a | AF700 | Biolegend |
| Anti-human CD56 | HCD56 | PE-Dazzle 594 | Biolegend |
| Anti-human CD8 | SK1 | Pe-Cy7 | Biolegend |
| Anti-human CD4 | SK3 | BV510 | Biolegend |
| Anti-human NKG2A | 131411 | BV421 | BD |
| Anti-human PD-1 | EH12.2H7 | PE | Biolegend |
| Anti-mouse B220 | RA3-6B2 | BUV395 | BD |
| Anti-mouse CD107a | 1D4B | BUV395 | BD |
|  |  | PE | Biolegend |
| Anti-mouse CD44 | IM7 | BUV395 | BD |
|  |  | Pacific Blue | Biolegend |
| Anti-mouse CD25 | PC61 | BUV395 | BD |
|  |  | BV650 | Biolegend |
|  |  | APC | Biolegend |
| Anti-mouse CD4 | RM4-4 | BUV496 | BD |
|  | RM4-5 | PerCp | Biolegend |
| Anti-mouse MHCII | M5/114.15.2 | BUV496 | BD |
|  |  | PE-Dazzle 594 | Biolegend |
|  |  | AF700 | Biolegend |
| Anti-mouse CD3 | 17A2 | BUV563 | BD |
|  |  | BUV737 | BD |
|  |  | BV650 | Biolegend |
|  |  | AF700 | Biolegend |
|  |  | PerCp Cy5.5 | Biolegend |
| Anti-mouse TCRδ | GL3 | BUV615 | BD |
| Anti-mouse CD11c | N418 | BUV615 | BD |
|  |  | BV750 | Biolegend |
|  |  | PerCp Cy5.5 | Biolegend |
| Anti-mouse PD-1 | 29F.1A12 | Pe-Cy7 | Biolegend |
|  |  | BV605 | Biolegend |
| Anti-mouse CD103 | 2E7 | BUV615 | BD |
|  |  | BV711 | Biolegend |
|  |  | BV785 | Biolegend |
|  |  | BV510 | Biolegend |
| Anti-mouse CD69 | H1.2F3 | BUV661 | BD |
|  |  | BV785 | Biolegend |
|  |  | PE-Dazzle 594 | Biolegend |
| Anti-mouse CD19 | 1D3 | BUV661 | BD |
|  | 6D5 | APC | Biolegend |
| Anti-mouse NK1.1 | PK136 | BUV737 | BD |
|  |  | BV421 | Biolegend |
| Anti-mouse TCRβ |  | BUV737 | BD |
| Anti-mouse CD45 | 30-F11 | BUV805 | BD |
| Anti-mouse F4/80 | BM8 | Pacific Blue | Biolegend |
| Anti-mouse FoxP3 | FJK-16s | eFluor 450 | Thermofisher Scientific |
| Anti-mouse NKG2A | 20D5 | Pacific Blue | BD |
|  | 16A11 | PE | Biolegend |
|  |  | Pe-Cy7 | Biolegend |
| Anti-mouse PD-L1 | MIH5 | BV421 | Biolegend |
| Anti-mouse PD-L1 | 10F.9G2 | PerCp Cy5.5 | Biolegend |
| Anti-mouse Ly6G | 1A8 | BV510 | Biolegend |
| Anti-mouse CD62L | MEL-14 | BV510 | Biolegend |
|  |  | Pe-Cy5 | Biolegend |
|  |  | PE-Dazzle 594 | Biolegend |
|  |  | PerCp Cy5.5 | Biolegend |
|  |  | AF700 | Biolegend |
| Anti-mouse CD8α | 53.-6.7 | BV570 | Biolegend |
|  |  | BV711 | Biolegend |
|  |  | BV785 | Biolegend |
| Anti-mouse CD27 | LG.3410 | BV650 | Biolegend |
|  |  | PE-Dazzle 594 | Biolegend |
|  |  | PerCp Cy5.5 | Biolegend |
|  | LG-7F9 | AF700 | Thermofisher Scientific |
| Anti-mouse TIM-3 | RMT3-23 | BV711 | Biolegend |
|  |  | APC | Biolegend |
| Anti-mouse ICOS | C398.4A | BV711 | Biolegend |
|  |  | BV750 | Biolegend |
| Anti-mouse CD11b | M1/70 | BV750 | Biolegend |
|  | HK1.4 | APC | Biolegend |
|  |  | AF700 | Biolegend |
|  |  | BV785 | Biolegend |
| Anti-mouse Qa-1b | 6F10.1A6 | PE | BD |
| Anti-mouse Ki67 | B56 | BV786 | BD |
|  | 16A8 | AF488 | Biolegend |
|  |  | PE | Biolegend |
|  |  | PE-Dazzle 594 | Biolegend |
| Anti-mouse NKG2D | CX5 | PE | Biolegend |
|  |  | PE-Dazzle 594 | Biolegend |
| Anti-mouse CD28 | 37.51 | PE | Biolegend |
|  |  | PE-Dazzle 594 | Biolegend |
|  |  | PerCp Cy5.5 | Biolegend |
| Anti-mouse Perforin | S16009B | PE | Biolegend |
| Anti-mouse Grzmb | GB11 | AF647 | Biolegend |
| Anti-mouse CD40L | SA047C3 | APC | Biolegend |
| Anti-mouse CD40 | 3/23 | Pe-Cy7 | Biolegend |
|  | 3/23 | PE-Dazzle 594 | Biolegend |
| Anti-mouse T-bet | 4B10 | Pe-Cy7 | Thermofisher Scientific |
| Anti-mouse EpCAM | G8.8 | FITC | Biolegend |
